# Four Potential Mechanisms Underlying Phytoplankton-Bacteria Interactions Assessed Using Experiments and Models

**DOI:** 10.1101/2025.01.03.631208

**Authors:** Osnat Weissberg, Dikla Aharonovich, Zhen Wu, Michael J. Follows, Daniel Sher

## Abstract

Phytoplankton growth and death depend on interactions with heterotrophic bacteria, yet the underlying mechanisms remain mostly unclear. We ask whether mathematical models explicitly representing four putative mechanisms of interaction (overflow metabolism, mixotrophy, exoenzymes and ROS detoxification) can recapitulate laboratory co-cultures between *Prochlorococcus* and 8 heterotrophic bacteria, ranging from synergism to antagonism. Two fundamentally distinct modes of interaction emerge from the models: i) Organic carbon and nitrogen recycling through exoenzymes or overflow metabolism, where high biomass of both organisms leads to more productive systems and more recalcitrant organic matter. ii) Detoxification of ROS, where a small number of "exploited" heterotrophs are sufficient to support *Prochlorococcus* survival. Cross feeding is likely more common in laboratory co-cultures, and most models cannot reproduce total inhibition of *Prochlorococcus*, suggesting additional mechanisms such as allelopathy. Key model parameters determining co-culture outcome are related to cell death and biomass recycling, providing priorities for future research.

---

Photosynthetic marine microbes (phytoplankton, both prokaryotes and eukaryotes) perform one-half of the primary production on Earth ^1^. The carbon (C) fixed by phytoplankton is released as they grow and die, feeding the entire food web, including heterotrophic bacteria, micrograzers, zooplankton, fish and ultimately human societies ^2,3^. Co-occurring marine microbes, including heterotrophic bacteria, affect the growth and death of phytoplankton (e.g. ^4–11^, potentially impacting ecosystem structure ^12–16^. Even though they occur on the scale of micrometers ^17,18^, these interactions may have far-reaching implications for global biogeochemical cycles, weather and the climate of our planet ^2^.

Phytoplankton-bacteria interactions can be highly complex and involve multiple mechanisms. Firstly, phytoplankton fix the dissolved organic carbon (DOC) utilized by heterotrophic bacteria, while at the same time both organisms compete for essential inorganic nutrients (recently reviewed by ^19–26^). As bacteria take up and utilize phytoplankton-derived DOC, they also modify it, changing its availability to other organisms, including the producing phytoplankton (e.g. ^27–29^). Secondly, bacteria may support phytoplankton growth by de-toxifying waste products, including reactive oxygen species (ROS, ^6,30–32^). Finally, phytoplankton and heterotrophic bacteria can directly signal to each other through specific infochemicals, or directly attack each other using allelochemicals or toxins (e.g. ^7,8,33–36^). Thus, phytoplankton-bacteria interactions encompass multiple mechanisms and result in a range of ecological interaction outcomes, from facilitation through commensalism to competition and allelopathy.

Given the diversity of phytoplankton and bacteria and the multiple potential mechanisms of interaction, it is difficult to determine which mechanism can underlie which interaction. In this study, we develop a modeling framework that mathematically describes the hypothesis that interactions between phytoplankton and heterotrophic bacteria are primarily metabolic, encompassing competition for nutrients, the release, modification, uptake and utilization of labile DOM, and the detoxification of ROS (Figure 1A). We then ask whether these models can recapitulate the results of laboratory co-cultures between *Prochlorococcus* MED4, representing a globally abundant clade of marine pico-cyanobacteria, and eight phylogenetically diverse heterotrophic bacteria (see below). If these relatively simple and tractable models are able to capture key features of the laboratory experiments, they could then be used to understand the effect of interactions on ecologically and biogeochemically important processes such as carbon fixation and DOC production. Cases where models fail to recapitulate experimental results suggest a structural deficiency in the model and the need to incorporate additional mechanisms, such as direct signaling or allelopathy.

**Figure 1.**
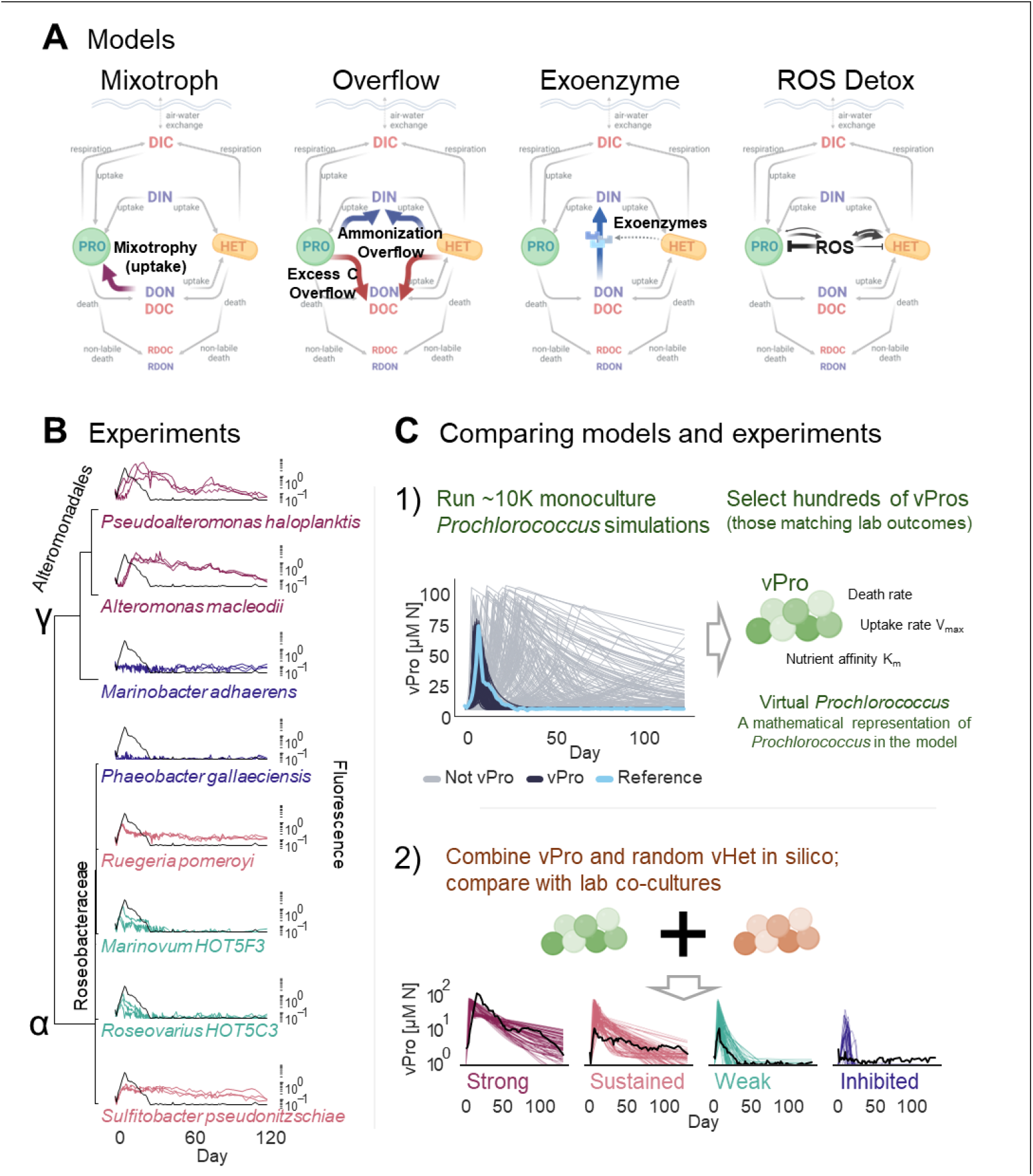
combining models and experiments to study co-culture mechanisms. A. Four models representing putative metabolic mechanisms underlying the interaction between the *Prochlorococcus* and the heterotroph. DIN, DIC, DON, DOC – dissolved inorganic and organic forms of nitrogen and carbon. RDON, RDOC – recalcitrant DON or DOC, PRO – *Prochlorococcus*, HET - heterotroph. B. Growth and decline curves f *Prochlorococcus* in coculture with 8 heterotrophic bacteria, measured using bulk fluorescence and organized based on the phylogeny of the heterotroph in co-culture. Y axes are logarithmic scale, colors inndicate coculture outcome, black line represents growth of *Prochlorococcus* axenic controls. α – alphaproteobacteria, γ – gammaproteobacteria. C. Outline of the modeling and computational analysis, where virtual *Prochlorococcus* (vPros) selected to match growth of axenic controls were simulated in cooculture with randomly selected virtual heterotrophs (vHets) and the resulting in-silico curve classified to determine which experimental outcome they are more similar to. Created in BioRender.com

Our model phytoplankton, *Prochlorococcus*, is a globally-abundant and ecologically-important clade of phytoplankton which have been extensively studied at the physiological, genomic and ecological-oceanographic levels (reviewed by ^37–39^). *Prochlorococcus* (and its close relative, *Synechococcus*) interact with heterotrophic bacteria in laboratory co-cultures, and that these interactions may differ based on the identity of the interacting organisms (e.g. ^6,40–45^). Due to their small size, *Prochlorococcus* have a very small phycosphere ^17^. Thus, while contact-mediated interactions cannot completely be ruled out, it is expected that most interactions are based on the exchange of dissolved molecules ^18,45,46^. As heterotrophic counterparts, we have selected eight strains of marine bacteria which have been studied intensively in the context of their interactions with diverse phytoplankton, including *Prochlorococcus, Synechococcus*, coccolithophores and diatoms, spanning the range from synergism to antagonism (Table 1, Supplementary Text 1, ^6,10,36,45,47–49^)). We focus on conditions of nitrogen (N) starvation, which are prevalent in the global ocean ^21^, and where microbial interactions are critical for the long-term survival of *Prochlorococcus* ^40,50^.

**Table 1:** Strains of heterotrophic bacteria co-cultured with *Prochlorococcus*.

| Group | Name | Reason Selected | Initial conc. cell ml<sup>-1</sup> | MPN on day 120 cell ml<sup>-1</sup> | Ref |
| --- | --- | --- | --- | --- | --- |
| **Strong** | *Alteromonas macleodii* HOT1A3 | Synergistic or inhibitory interaction depending on *Prochlorococcus* strain and dose | 1e6 | 5e7-5e10 | <sup>48</sup> |
| **Strong** | *Pseudoalteromonas haloplanktis* CIP 108707 | FBA model, chemotaxis to *Prochlorococcus* extracellular products. | 1e7 | 5e6-5e7 | <sup>49</sup> |
| **Sustained** | *Ruegeria pomeroyi* DSS-3 | Genetically tractable model organism, survives long term starvation in co-culture with *Synechococcus* | 1e7 | 0 | <sup>6</sup> |
| **Sustained** | *Sulfitobacter pseudonitzschiae* SMR1 | Interactions with diatoms, including production of plant hormone IAA. | 1e7 | 5e5-5e6 | <sup>10</sup> |
| **Weak** | *Marinovum* HOT5F3 | negative interaction with *Prochlorococcus* | 1e7 | 5e4 | <sup>45</sup> |
| **Weak** | *Roseovarius* HOT5C3 | Positive/neutral interaction with *Prochlorococcus* | 1e7 | 5e2 | <sup>45</sup> |
| Inhibited | *Phaeobacter gallaeciensis* | Remarkable surface (abiotic/biotic) colonizer, kill diatoms and other bacteria. Promotes growth than kills *Emiliania huxleyi* | 2.5e6 | 5e7-5e10 | <sup>36</sup> |
| Inhibited | *Marinobacter adhaerens* HP15 | Affects diatom aggregation and growth in co-culture | 1e7 | 5e6-5e7 | <sup>47</sup> |

## Results

### Laboratory co-cultures show diverse outcomes

The growth and death dynamics of *Prochlorococcus* MED4 in co-culture with the eight heterotrophic bacteria could be divided into four main groups based on the maximum fluorescence and the decline stages of the curves (Figure 1B). *Alteromonas macleodii* and *Pseudoalteromonas haloplanktis* reached either similar or higher fluorescence than the axenic controls, and declined much more slowly, consistent with previous studies of diverse *Alteromonas* strains ^40,48,51^. We defined this outcome as "strong". *Ruegeria pomeroyi* and *Sulfitobacter pseudonitzschiae* both reached a lower maximal fluorescence than the axenic control but then maintained higher fluorescence for the entire 129 days of the experiment, which we defined as a "sustained" outcome. *Marinovum HOT5F3* and *Roseovarius HOT5C3* both grew to a lower fluorescence than the axenic controls and declined as rapidly as them, with long-term fluorescence below the limit of detection (defined as "weak"). Finally, no *Prochlorococcus* growth was observed in co-culture with *Marinobacter adhaerens HP15* and *Phaobacter gallaciensis*, and we defined this outcome as "inhibited".

Some of the results of the *Prochlorococcus*-heterotroph co-cultures are consistent with the effect of the same heterotrophs on different phytoplankton, whereas others differed. For example, *Ruegeria pomeroyi* and *Sulfitobacter pseudonitzschiae*, both of which produced a sustained outcome, support the growth of *Synechococcus* and diatoms, respectively, through the remineralization of organic carbon sources and the production of ammonium ^6,10^. Similarly, *Phaeobacter gallaeciensis* which inhibits *Prochlorococcus*, produces antibiotics and algaecides in coculture with diatoms, in a manner which depends on the diatom growth stage ^36^. In other cases, the outcome is different, for example, *Marinobacter adhaerens HP15* which inhibits *Prochlorococcus* has more complex interactions with diatoms, adhering to them, affecting the exudation of transparent exopolymers and inducing aggregation, but not inhibiting them ^47^.

### Models of different interaction mechanisms differ in their ability to recapitulate lab co-cultures

We next developed four mathematical models of phytoplankton and heterotrophic bacteria growing on and exchanging inorganic and organic forms of carbon (C) and Nitrogen (N), asking to what extent each model can represent the laboratory co-cultures. Each model represents a putative metabolic mechanism that could underlie the interactions between the two organisms (Figure 1A): 1) Mixotrophy, where phytoplankton can utilize organic C and N (DOC and DON) in addition to inorganic forms ^52–54^; 2) Overflow metabolism, where organisms take up excess C or N, but cannot use it to build new biomass due to limitation of the other element. The excess C or N are modified and released through exudation ^55^; 3) Exoenzymes which degrade extracellular DON to DIN (DIN represents also simple organic N sources such as amino acids that can be utilized by both organisms) ^56,57^; 4) Detoxification of Reactive Oxygen Species (ROS) by the heterotrophic bacteria ^30–32^. All four models have a common, basic structure, where phytoplankton support heterotrophic bacteria through photosynthesis and the release of DOC and DON during cell death or, in our idealized model, passive leakage. Both organisms compete for DIN. Within each organism, C and N are taken up into internal stores, and subsequently combined into biomass, enabling the model to capture varying elemental stoichiometry ^52,58^. Part of the DOC and DON released during cell death is in a form that is not bioavailable, representing the biogeochemically important process of the production of recalcitrant DOM ^59^. For a more detailed description of the model structures please see Supplementary text 2.

We have previously shown that there are multiple combinations of different model parameters (e.g. K^m^, V_max_, mortality and exudation rates), which can equally well fit experimental data ^60^. Therefore, rather than choosing a single set of parameters to represent *Prochlorococcus*, we performed a Monte Carlo simulation to generate 10,000 different parameter combinations, selecting from these a subset of 30-55 "virtual *Prochlorococcus*" (vPros) which best fit the axenic growth data (Figure 1C, Supplementary Texts 3-5). We then simulated the growth of these vPros, which can be thought of as different strains of *Prochlorococcus*, with thousands of randomly generated "virtual Heterotrophs" (vHets, Figure 1C). Finally, we classified each of the "in-silico" vPro-vHet combinations as one of the growth phenotypes described above, using Machine Learning, and filtered the results to retain 2,216-6,313 vPro-vHet pairs where the C:N ratio of the two interacting organisms was feasible biologically (3.5-10, ^60^, Supplementary Text 5).

From the four modelled mechanisms (mixotrophy, overflow, exoenzymes and ROS detoxification) three were able to recapitulate all of the lab outcomes, albeit at different ratios (Figure 2A, B, Supplementary Figure S12A). In contrast, the mixotrophy model, as well as the basic model without any of the mechanisms, were only able to recapitulate two outcomes – weak and neutral. Similar results were obtained in a second round of simulations where the number of vPros was increased by perturbing the parameters of the existing vPros from the first run (Supplementary Figure S12).

**Figure 2.**
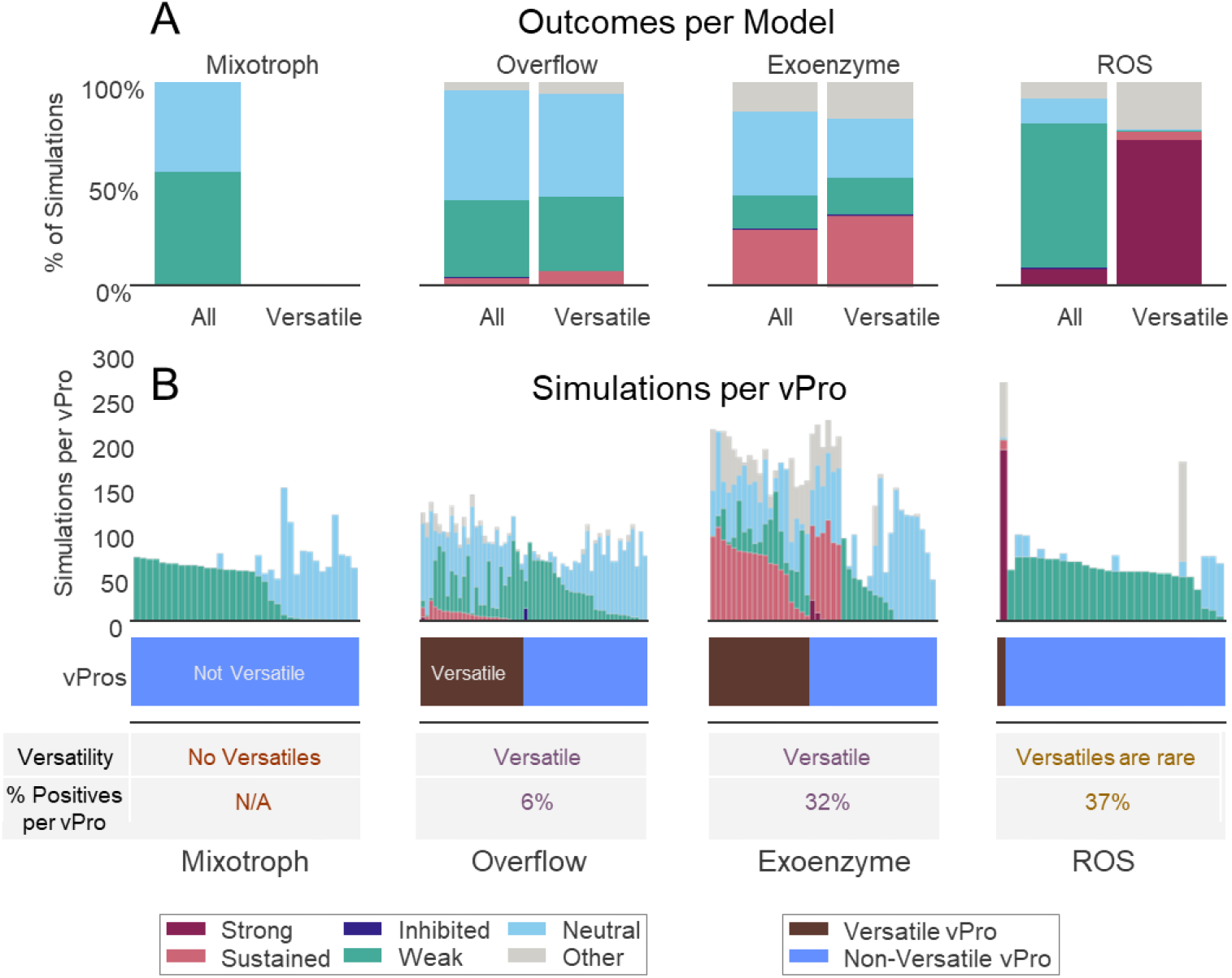
Outcome of the simulations. A. Simulation outcomes in each model when either all vPro-vHet pairs are considered, or only versatile vPros. B. Simulation per vPro, each bar corresponds to simulations of a specific vPro with multiple vHets. vPro versatility is indicated at the bottom of the panel.

We next focused on the subset of vPros that were able to recapitulate both positive (strong and/or sustained) and negative (weak and/or inhibited) outcomes, because in lab co-cultures a single *Prochlorococcus* strain exhibited different outcomes with different heterotrophs (Figure 1B, Figure 2B, Supplementary Figure S12B). There were many more such "versatile" vPros in the overflow and exoenzyme models, with only one in the ROS model and none in the mixotroph. Additionally, the versatile vPros in the exoenzyme and overflow models produced mostly sustained outcomes, whereas those in the ROS model reproduced primarily strong outcomes. There were very few inhibitory outcomes in the models (0-0.3%), none of which reproduced the complete lack of *Prochlorococcus* growth observed in laboratory cultures (Supplementary Figure S13).

These results suggest that within the range of the model parameters tested, the exoenzyme model is best at capturing the variability of lab co-culture outcomes, although the overflow model produced a similar fraction of versatile vPros, and the ROS model was better at capturing the strong outcome. Each of these mechanisms has been invoked to explain the dynamics of phytoplankton-bacteria interactions (e.g. ^6,31^, yet our results suggest that mixotrophy, despite its ecological importance ^52^, is less likely to be a single mechanism underlying the *Prochlorococcus*-heterotroph interactions observed in the lab. We therefore focused on the other three models for subsequent analyses.

### Loss processes are important determinants of *Prochlorococcus* versatility and the outcome of the co-cultures

We next asked which *Prochlorococcus* and heterotroph traits determine interaction outcome. These traits are encoded in the mathematical models as the values of the parameters describing processes such as nutrient uptake, mortality, and DOC/DON modification and recycling. V_max_ and K_m_ for uptake were only weakly correlated with the outcome of the co-cultures, despite these parameters often considered fundamental for understanding ecosystem dynamics (Figure 3A, Supplementary Figures S5A, S6A, ^61–63^). Instead, co-culture outcome was strongly and negatively correlated with the mortality of both interacting organisms, as well as to the fraction of released biomass bioavailable to the other organism (termed γ in the models, Figure 3B, Supplementary Figures S5B, S6A). Mortality is described in three of the models (mixotrophy, overflow and exoenzyme) as a single, intrinsic, exponential mortality value, whose value for *Prochlorococcus* needs to be high for the models to fit the rapid decline observed in axenic cultures and weak co-cultures (0.46 ± 0.17 day^−1^, in agreement with ^48^). In contrast, in the ROS model, mortality is the sum of intrinsic and ROS-depended mortality (Supplementary Figure S1, equations 25,26 in Supplementary Text 2). For a vPro in the ROS model to be versatile it needs to have very low intrinsic mortality rates (0.02 ± 0.01 day^−1^), coupled with high ROS production rate, low ROS degradation rate and high ROS toxicity (Supplementary Figures S5C, S6C). In these vPros heterotrophic bacteria can elicit different co-culture outcomes through variations in ROS production and degradation (Figure 3C, Supplementary Text 5).

**Figure 3.**
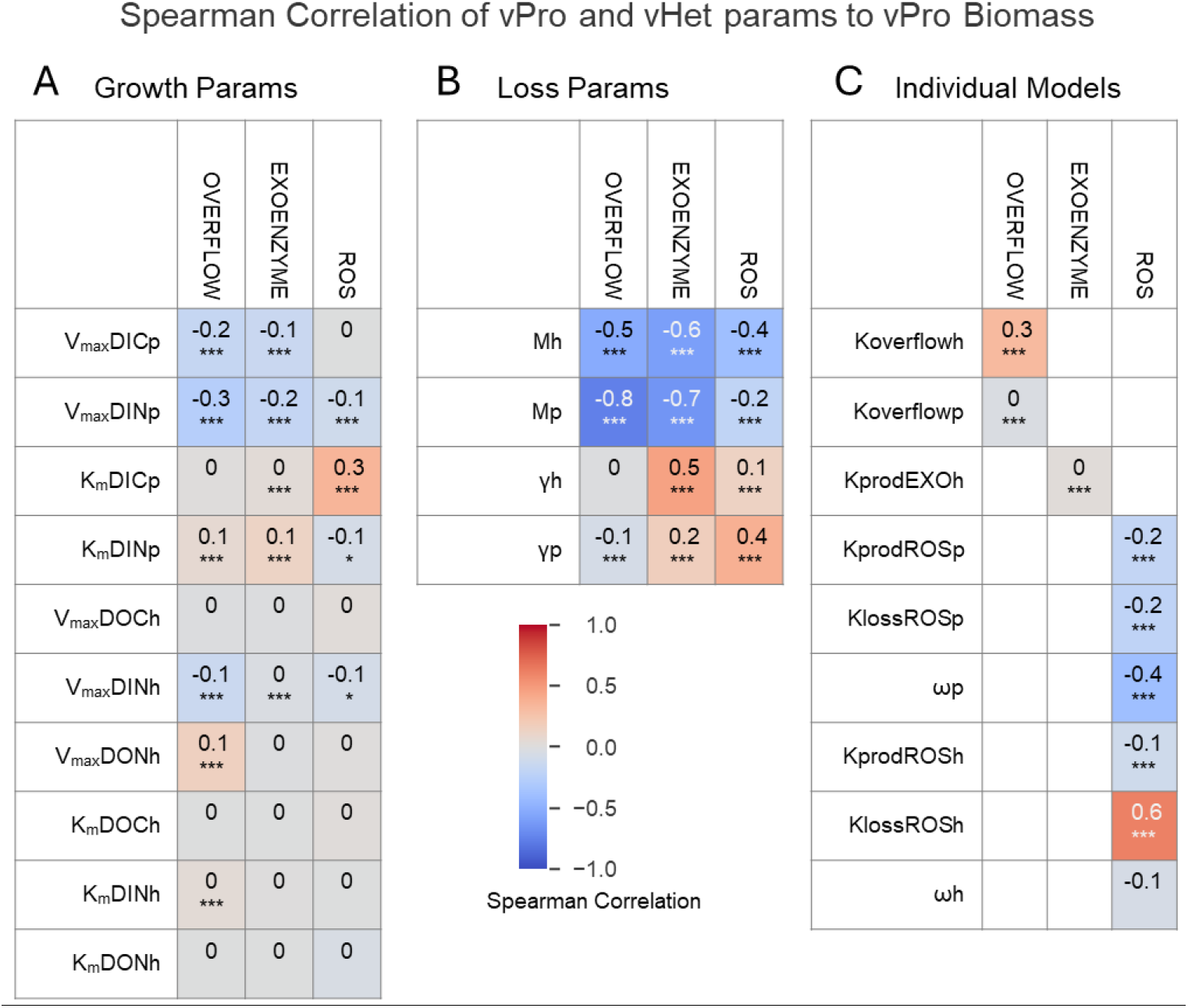
Spearman correlation between model parameters and *Prochlorococcus* biomass, which is associated with co-culture outcome (Supplementary Figure S14). A) Loss parameters B) Growth parameters C) Parameters of individual models. */**/*** - corrected p-value of the correlation (0.05, 0.01 and 0.001, respectively). Mp/Mh: death rates of vPro and vHet respectively, γp/γh: Rates of dead matter lability, V_max_DICp/V_max_DINp/V_max_DOCh/ V_max_DINh/V_max_DONh maximum uptake rates for DIC, DIN, DOC, DIN and DON respectively, K_m_DICp/K_m_DINp/K_m_DOCh/K_m_DINh/K_m_DONh nutrients affinities for DIC, DIN, DOC, DIN and DON respectively. Koverflowh/Koverflowp: maximum overflow rates, KprodEXOh: rate of exoenzyme degradation, KprodROSp/KprodROSp: ROS production rate, KlossROSp/ KlossROSp: ROS breakdown rate, ωp/ωh: ROS toxicity

In the recycling-based models (overflow and exoenzyme), regardless of the specific outcome, the biomass of the vHet was higher than that of the vPro, and they were highly correlated (Pearson correlation 0.77-0.86, Figure 4A, B). During the extended culture decline stage, the heterotrophs were mostly C limited, while *Prochlorococcus* was N limited (Figure 4D). Non-intuitively, *Prochlorococcus* biomass was highly correlated with the fluxes of DON to DIN through the overflow and exoenzyme systems (Figure 4C), but only moderately correlated with the rate constants of these processes (Figure 3C). This suggests a tightly coupled system where the growth of each organism depends on that of the other, and where relatively high biomass and high bioavailability of released organic matter (high γ, equations 5, 6, 10, 11 in Supplementary Text 2, Supplementary Figures S5B, S6A) sustain large fluxes of C and N between the two organisms despite low mortality. Paradoxically, these systems require a mechanism for DON recycling (overflow or exoenzymes), but the efficiency of these mechanisms is less important in determining the outcome of the interactions.

**Figure 4.**
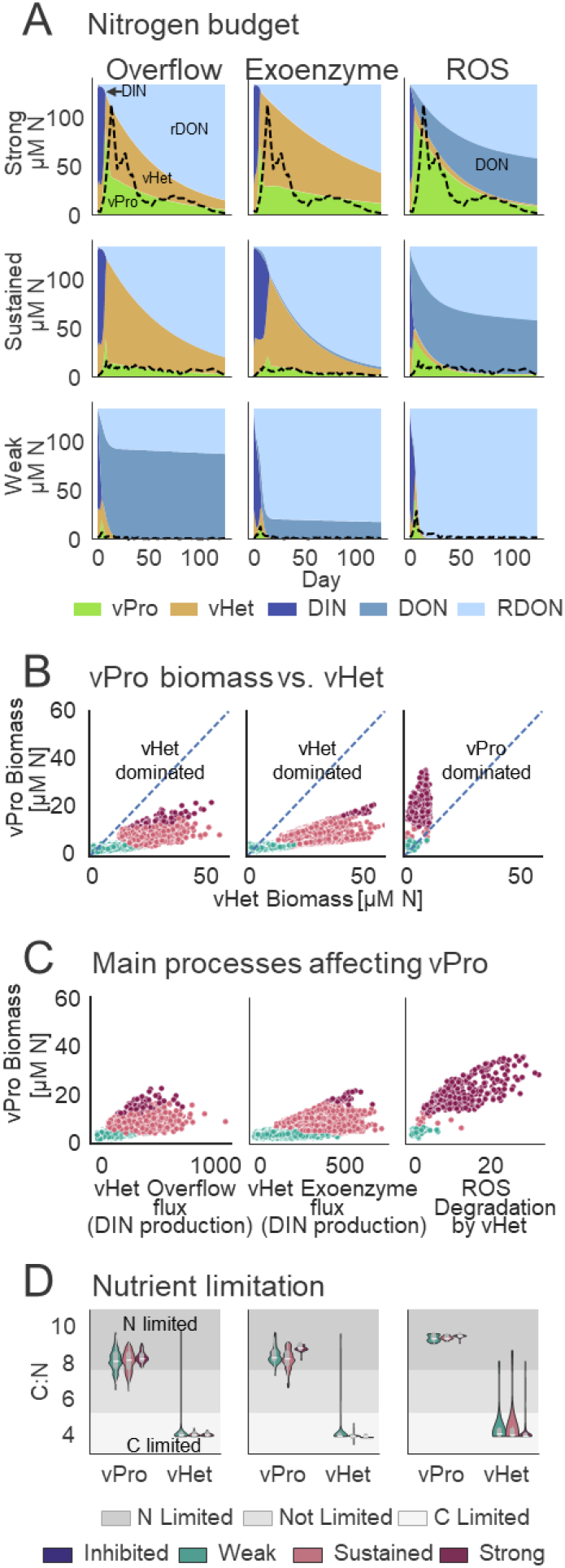
Details of modeled co culture outcomes. A) Breakdown of the nitrogen budget over time, the simulation most similar to the experimental results is shown for each outcome/model. B) mean N biomass of vPro vs. mean N biomass of vHet. Pearson correlations: Overflow 0.86, Exoenzyme 0.77, ROS 0.45. C) C: Main processes affecting outcome. D) C:N ratios during extended decline stage.

The ROS model predicted a different system, whose biomass was dominated by *Prochlorococcus* (Figure 4A, B). In this model γ is not constrained, suggesting a secondary importance of nutrient remineralization compared to ROS detoxification in this model. The heterotrophs were C limited, grew very slowly, yet in the strong and sustained outcomes produced just enough biomass to provide the ROS detoxification "ecosystem service" needed to enable vPro growth (Figure 4C). At the same time, the lack of a mechanism to recycle DON to DIN led to an accumulation of DON and to N limitation of *Prochlorococcus* (Figure 4A, D). In the weak and inhibited outcomes, either the heterotrophs were not able to grow (e.g. high death rate and/or low V_max_ for organic C) or could not provide sufficient ROS detoxification (e.g. low detoxification rate). In both cases this led to the collapse of both interacting organisms (e.g. figure 4A, right bottom panel). Thus, even though each organism in the co-culture depended on a resource provided by the other for growth and survival, these systems were more "exploiting" than the synergistic systems predicted by the recycling-based models ^64^.

While not all of the model predictions can be tested with our experimental data, flow cytometry and estimates of culturable heterotrophic bacteria during long term co-cultures both suggest that most of the cells are heterotrophs (Table 1, Supplementary Text 7, Supplementary Figure S11C, D). This is more reminiscent of the predictions of the overflow or exoenzyme models. Therefore, we propose that for most of the heterotrophs tested recycling of N (and possibly C) is the main mechanism underpinning the *Prochlorococcus*-heterotroph interactions.

### Does combining multiple models increase biological realism?

The models describing each of the four interaction mechanisms are relatively simple and tractable, yet in nature it is likely that these mechanisms occur together. Combining multiple models did not increase two measures of model quality (the fraction of versatile vPros and model-data fit probability) compared with models of a single mechanism (Supplementary Figures S15, S16). Moreover, the outcome of combined models was highly consistent with that of one of the individual models included in the combination. Specifically, a Strong outcome was associated with the ROS mechanism, although the fraction of Strong outcomes was reduced when combined with most other models. Adding the ROS mechanisms to any other model resulted in most of the Strong outcomes having higher *Prochlorococcus* than heterotroph biomass (Supplementary Figure S16A), which contrasts with the laboratory results. Therefore, while the ROS detoxification mechanism may strongly affect model results, it is likely that in the lab co-cultures recycling is the dominant process.

The Sustained outcome was associated mainly with the exoenzyme model, whereas the Weak model was mainly associated with the overflow model. The fraction of Weak outcomes was reduced when the overflow and exoenzyme models were combined, hinting at the importance of extracellular degradation compared with overflow metabolism in long-term survival (Supplementary Figure S16A).

Surprisingly, the mixotrophy model increased the fraction of inhibited outcomes when combined with the exoenzyme and/or overflow models, and these model combinations were able to recapitulate the complete lack of *Prochlorococcus* growth observed in some laboratory co-cultures (EM, OM and EOM models in Supplementary Figure S13). Since the inhibited vPros had low photosynthesis rate (low V_max_DICp), and a high uptake for DOC, it is possible that the inhibition was due to competition for DOC (Supplementary Figure S17). Nevertheless, even in these simulations the overall fraction of inhibited vPro-vHet interactions was small (<2.5%, Supplementary Figure S16), reinforcing our hypothesis that full inhibition of *Prochlorococcus* is due to additional models such as allelopathy.

### Biogeochemical implications

The experiments and models presented here describe highly simplified systems, differing fundamentally from the complex and dynamic ocean. Nevertheless, it is tempting to ask to what extent the mechanisms studied here could affect higher-order processes such as primary production, carbon and nitrogen use efficiency, and carbon sequestration. The net community production of the system, defined here as the amount of organic C remaining at the end of the simulation (in organismal functional biomass and C storage, DOC and recalcitrant DOC), was highest in the Strong and Sustained outcomes of the two recycling models (Figure 5A). The carbon use efficiency (CUE, a measure of the capacity of a given system to effectively retain carbon), was lower in the Strong and Sustained outcomes, where both organisms benefited from the interaction (Figure 5B). This is because the Strong and Sustained interactions were defined by *Prochlorococcus* surviving much longer, maintaining a larger heterotrophic population (Figure 4A, B). During this extended culture decline stage, maintenance rather than growth dominates, and most of the carbon is respired and not integrated into biomass. Nevertheless, despite the lower CUE, the net productivity of the Strong and Sustained co-cultures is 5-10 times higher than that of the Weak and Inhibited ones (Figure 5A).

**Figure 5.**
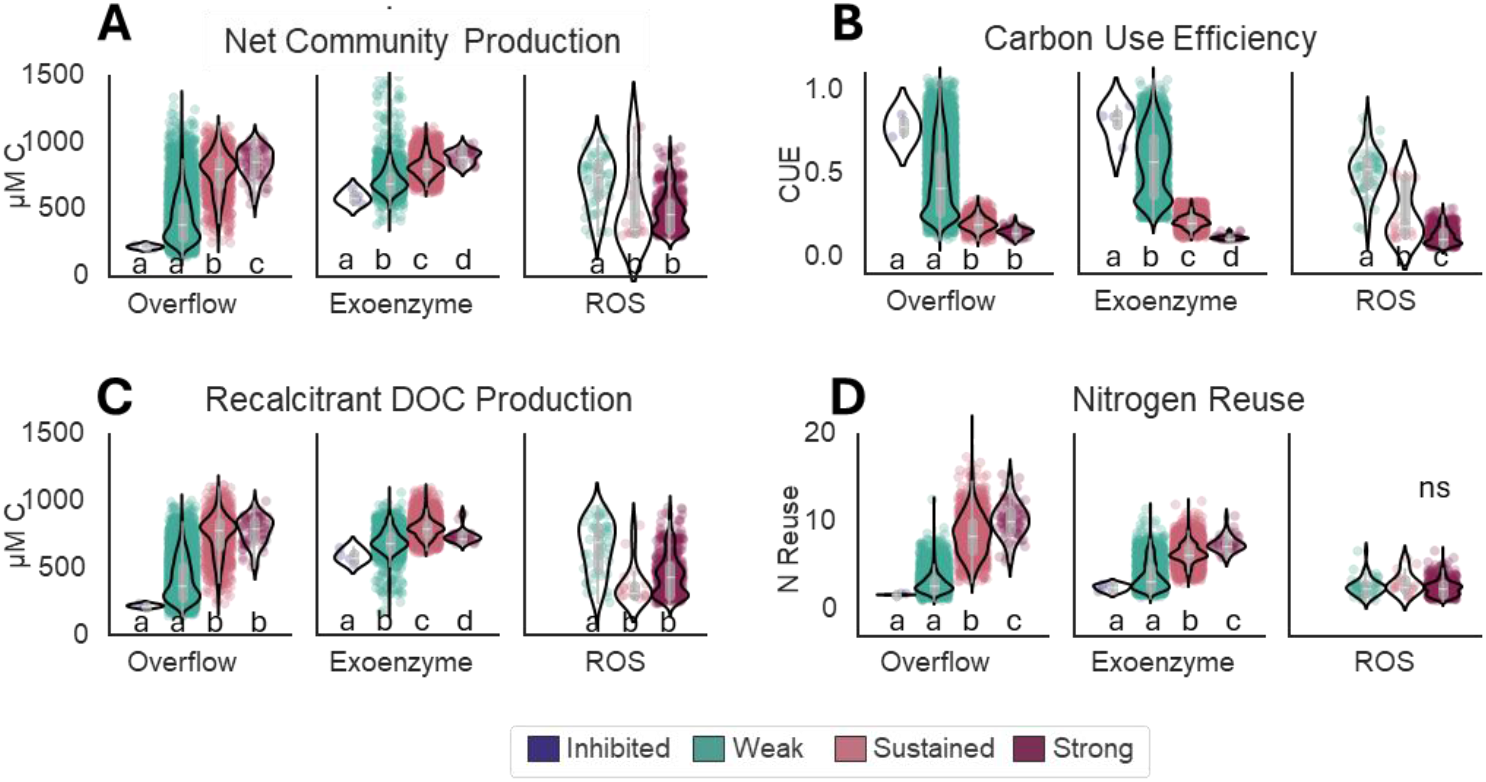
Biogeochemical traits. A) Organic Carbon at the end of the experiment, including DOC, RDOC and biomass. B) Carbon use efficiency. C) Recalcitrant DOC (RDOC) at the end of the experiment. D) Nitrogen reuse. Black outline shows violin plot describing the distribution of the simulations, inside grey box shows the quartiles, colored dots represent individual simulations. Letters indicate grouping based on Anova and Tukey post hoc.

N reuse (the number of times a nitrogen atom was incorporated into biomass) was highest in the Strong and Sustained phenotypes of the recycling models, reflecting the metabolic coupling between the phototrophs and the heterotrophs (Figure 5D). This may be reminiscent of the rapid and significant N recycling in the photic zone of oligotrophic oceans (e.g. low new production), where *Prochlorococcus* are the most dominant phytoplankton ^65,66^. Finally, most of the C remaining in the system was in recalcitrant form (rDOC, Figure 5C). Carbon export and sequestration in the oceans is mostly driven by particle sinking, yet DOC export is a major process, responsible for up to 20% of the carbon export globally, and ∼50% of export in oligotrophic areas of the ocean where *Prochlorococcus* is the dominant phytoplankton ^67^. Most ocean models underestimate DOC production and export in the oligotrophic gyres ^58^, and our results suggest that these processes would be preferred when the phytoplankton and heterotrophic bacteria are strongly coupled through remineralization processes such as the overflow and exoenzymes mentioned above.

## Discussion

The intricate network of interactions between phytoplankton and heterotrophic bacteria is ecologically and biogeochemically important. Our results show that models explicitly representing DON recycling (though metabolic overflow and/or exoenzymes) are necessary and sufficient to recreate most of the diverse outcomes observed in experimental laboratory co-cultures. These mechanisms are therefore good candidates for more detailed study, and for incorporation into more complex ecosystem or Earth System models ^68^, especially with regard to the oligotrophic nutrient-poor ocean, where recycling of the limited available nutrients likely plays an important role. In contrast, while ROS detoxification enables *Prochlorococcus* to survive multiple types of stress, including temperature extremes and extended darkness ^30–32,69^, our results suggest that this mechanism is less likely to be the key mechanism underlying long term nitrogen starvation. Nevertheless, ROS detoxification may strongly impact the geographical abundance of *Prochlorococcus* ^30^. A more detailed ROS budget of the oceans will help understand when and where this mechanism is relevant for biogeochemical models. Mixotrophy provides an important source of carbon and energy for *Prochlorococcus* living at the bottom of the photic zone ^52^, and may also contribute significantly to the N budget of natural populations ^57^. Yet, this model does not recapitulate long term survival, despite providing *Prochlorococcus* access to the limiting resource, nitrogen. Unlike the other three mechanisms, which all describe different forms of cooperation, the mixotrophy model revolves around competition for both organic and inorganic resources. An interesting open question is why such competition alone was not able to recapitulate the complete inhibition of *Prochlorococcus* yet could do so when coupled with nutrient recycling. Regardless, very few vPro-vHet pairs resulted in the complete inhibition of *Prochlorococcus*, despite this outcome being relatively common in laboratory co-cultures (Figure 1B). We speculate that in such interactions other mechanisms may be involved, e.g. allelopathy mediated by toxins or antibiotics.

## Acknowledgements

This study was funded by the Israel Science Foundation (grant number 1786/20 to DS) and by the National Science Foundation - United States-Israel Binational Science Foundation (NSFOCE-BSF 1635070 and NSF-BSF 2246707 to DS). MJF is grateful for support from the Simons Foundation (CBIOMES, grant number 549931 to MJF).

## Code availability

The codes to simulate the different mechanistic models and analyze the simulation results are available at Github (https://github.com/wosnat/recycle_model/tree/main/STORE_model) under MIT license.

## Materials and Methods

### Coculture growth experiments

*Prochlorococcus* MED4 was grown in triplicate batch coculture with 8 different bacteria strains (Table 1) for 129 days under nitrogen stress conditions (pro99 low N). Triplicate batch cultures of axenic *Prochlorococcus* MED4 were used as negative control. The viability of *Prochlorococcus* in the co-cultures was checked by transferring to fresh media on days 42, 60, 81, 129. The resulting growth curves are clustered into 4 different interaction phenotypes via visual inspection of *Prochlorococcus* growth and decline curves.

*Prochlorococcus* growth was measured by auto fluorescence, a proxy for *Prochlorococcus* cell numbers during exponential growth and decline ^48^. Flow cytometry was run on the inoculated culture on days 20, 42, 60, 81, 129. There is a linear correlation between the fluorescence and the cell counts (R^2^=0.733, Pval=6.74e-42, supplementary Figure S11B). We used linear regression to convert the culture fluorescence to cell counts, and then converted these cell counts to culture biomass by multiplying by *Prochlorococcus* cell mass previously measured in our lab ^70^. We used these biomass approximations as basis for tuning our model.

The viability of the heterotrophs cultures was measured using the Most Probable Number (MPN) method, where serial 10-fold dilutions were inoculated into 96 well plates containing Marine broth media on day 129, and the growth of heterotrophs (well turbidity) assessed over 7 days.

### Strains and experiment set up

Axenic *Prochlorococcus* MED4 was maintained under constant cold while light (27 μmole photons m^−2^ s^−1^) at 22 °C ^45,71^. We used Pro99 media (natural seawater-based) that was modified by reducing the concentration of NH_4_^+^ from 800 μM to 100 μM (Pro99-LowN), resulting in *Prochlorococcus* entering stationary stage due to the depletion of available inorganic N ^60^. Heterotrophic bacteria (Table 1) were maintained in ProMM ^72^. At the start of each co-culture experiment, heterotroph cells from stationary-stage cultures (24-72 hour old) were centrifuged (10 minutes, room temperature, 10,000 g), the growth media decanted, and the cells re-suspended in the experimental media (Pro99-LowN). The *Prochlorococcus* cells (from mid-exponential cultures) and the re-suspended heterotroph cells were counted using BD FACSCanto II Flow Cytometry Analyzer Systems (BD Biosciences), and co-inoculated at initial concentrations of 1×10^6^ *Prochlorococcus* cells ml^−1^ and/or 1×10^7^ heterotroph cells ml^−1^. The experiment was performed using triplicate 20 ml cultures in borosilicate test tubes (2.5 cm diameter, 15 cm length).

### Fluorescence and Flow cytometry

Bulk chlorophyll fluorescence (FL) (ex440; em680) was measured approximately every 2 days using a Fluorescence Spectrophotometer (Cary Eclipse, Varian). Samples for flow cytometry were taken after 42, 60, 81, and 129 days, fixed with glutaraldehyde (0.125% final concentration), incubated in the dark for 10 min and stored in −80 °C until analysis. Fluorescent beads (2 μm diameter, Polysciences, Warminster, PA, USA) were added before flow cytometry as an internal standard. Data was acquired and processed with FlowJo software. Flow cytometry was performed unstained to count *Prochlorococcus* cells followed by staining with SYBR Green I (Molecular Probes/ ThermoFisher) to count heterotroph cells.

### Translating experimental data to model units

The model describes *Prochlorococcus* and heterotroph biomass in units of μmol N L^−1^. The fluorescence measurements were translated into biomass by first fitting fluorescence to cell concentration from flow cytometry (cells ml^−1^) using an OLS model (ordinary least squares, python package statsmodels), F_1,139_ = 385.2, p-value = 6.74e-42, adjusted R^2^ = 0.733. The cell concentration was then translated to μmol N L^−1^ by multiplying the number of cells by the *Prochlorococcus* N biomass of 8.9e-10 μmol N cell^−1^ and by 1000 to translate from ml to L. Measurements below limit of detection (<0.04 fluorescence) were filtered out.

| OLS Regression Results |  |  |  |  |  |  |
| --- | --- | --- | --- | --- | --- | --- |
| ===== |  |  |  |  |  |  |
| Dep. Variable: | FCM | R-squared (uncentered): | 0.735 |  |  |  |
| Model: | OLS | Adj. R-squared (uncentered): | 0.733 |  |  |  |
| Method: | Least Squares | F-statistic: | 385.2 |  |  |  |
| Date: | Sun, 29 Sep 2024 | Prob (F-statistic): | 6.74e-42 |  |  |  |
| Time: | 09:31:13 | Log-Likelihood: | -2500.8 |  |  |  |
| No. Observations: | 140 | AIC: | 5004. |  |  |  |
| Df Residuals: | 139 | BIC: | 5007. |  |  |  |
| Df Model: | 1 |  |  |  |  |  |
| Covariance Type: | nonrobust |  |  |  |  |  |
| ===== |  |  |  |  |  |  |
|  | coef | std err | t | P>\|t\| | [0.025 | 0.975] |
| ----- |  |  |  |  |  |  |
| FL | 2.626e+07 | 1.34e+06 | 19.627 | 0.000 | 2.36e+07 | 2.89e+07 |
| ===== |  |  |  |  |  |  |
| Omnibus: | 164.439 | Durbin-Watson: | 0.874 |  |  |  |
| Prob(Omnibus): | 0.000 | Jarque-Bera (JB): | 4684.735 |  |  |  |
| Skew: | 4.377 | Prob(JB): | 0.00 |  |  |  |
| Kurtosis: | 29.953 | Cond. No. | 1.00 |  |  |  |
| ===== |  |  |  |  |  |  |
| Notes: |  |  |  |  |  |  |
| [1] R<sup>2</sup> is computed without centering (uncentered) since the model does not contain a constant. |  |  |  |  |  |  |
| [2] Standard Errors assume that the covariance matrix of the errors is correctly specified. |  |  |  |  |  |  |

### Measurement of Kg – air/water exchange rate

As part of the model parameterization, we measured the gas exchange rate between sea water and air. We measured oxygen exchange as a proxy for CO_2_. We used 20ml of acid washed and autoclaved seawater in borosilicate test tubes, bubbled with nitrogen to deplete the oxygen in the media and measured the oxygen level for 24-48 hours using pyroscience sensor. The measurements were fitted using the formula: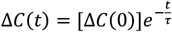, and then K_g_ was computed using the formula: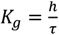.

### Mathematical Model Implementation

The model was implemented in python using differential equations in numpy and numba. Simulations were performed with solve_ivp (scipy) using BDF method and max_step=1000. The fit to experimental results was checked by geometric mean of the Root mean square error (RMSE) for N and C total biomass (i.e., the sum of functional biomass and respective store was compared to experimental N and C).

### Montecarlo simulation with random parameters

When simulating with random parameters, most of the model parameters were assigned predetermined values based on literature (see Supplementary text 3, Supplementary table S2), 3 or more of the parameters were assigned a value taken from a log uniform distribution of values, one order of magnitude below and one above the literature value.

### Selection of virtual *Prochlorococcus* (vPro)

Each vPro candidate was simulated in standalone mode on 2 media, PRO99-lowN and PRO99, and compared to experimental measurements. with a simulation defined as a vPro if the geometric mean of the RMSE of both N and C *Prochlorococcus* biomass on both media was less than 60.

The initial selection of vPro was based on 10,000 random Monte Carlo simulations. This resulted in the identification of 30-55 vPros per model. We then enlarged the sample size by running an additional 100 simulations per vPro, randomly changing 3-5 of its parameters, resulting in a final list of 300-700 vPros per model.

### Coculture simulations (vPro + vHet)

vPros were simulated in coculture with randomly selected heterotroph parameter values. We ran 10,000 simulations, using the vPros selected and varying the heterotroph parameters by 2 orders of magnitude.

We filtered out 1% of the simulations where there were technical problems (could not simulate for the entire span or overshot into negative biomass or nutrient pools). We filtered out additional ∼50% of the simulations where the heterotroph C:N ratio reached biologically infeasible values (below 3.5 C:N). Samples were classified into outcomes using a stacked logistic regression model based on comparison to the experimental N and C biomasses are well as computed features (see Extended Methods, Supplementary Text 6). Versatile vPros were defined as the subset of the vPro whose simulations reproduced both positive (Strong or Sustained) and negative (Weak or inhibited) outcomes.

### Analysis of the results

Productivity of the simulation was measured by mean total biomass. Total biomass (the sum of functional biomass and the respective store) was integrated over the length of the simulation using simpson integration (scipy) and divided by simulation length.

Fluxes in the model, such as nutrient uptake, recycling of dead cells, ROS production and degradation, are an additional output of the model simulation. Total integrated flux was computed by integration over the length of the simulation using simpson integration (scipy).

The correlation between model parameters and simulation outcome was measured by spearman correlation between mean total biomass and parameter values. We selected non-parametric spearman because parameter distributions were not normal due to the randomization method.

Nutrient limitation was measured as mean C:N during extended decline phase. N / C limitations were defined as C:N values where the regulation of uptake was less than 0.4 (equation 15 in Supplementary Text 2). Extended decline phase was defined as the duration of the simulation where C:N ratios of both organisms were stable, i.e., standard deviation of the C:N was less than 0.02. In most cultures, the extended decline phase lasted more than 50 days of the simulation. Final RDOC is the value of RDOC at the end of the simulation. Final organic C is the sum at the end of the simulation of DOC, RDOC, C stores and biomass for both organisms. Carbon use efficiency (CUE) is defined as the ratio between net primary productivity (NPP), i.e., final organic C, and gross primary productivity (GPP), measured as the total DIC uptake by *Prochlorococcus*. N reuse is measured by the ratio of integrated N uptake to initial N availability.

## References

1. Field, C. B., Behrenfeld, M. J., Randerson, J. T. & Falkowski, P. Primary Production of the Biosphere: Integrating Terrestrial and Oceanic Components. Science (1979) 281, 237–240 (1998).

2. Azam, F. & Malfatti, F. Microbial structuring of marine ecosystems. Nat Rev Microbiol 5, 966–U23 (2007).

3. Karl, D. M. Microbial oceanography: paradigms, processes and promise. Nat Rev Microbiol 5, 759–769 (2007).

4. Arandia-Gorostidi, N., Weber, P. K., Alonso-Sáez, L., Morán, X. A. G. & Mayali, X. Elevated temperature increases carbon and nitrogen fluxes between phytoplankton and heterotrophic bacteria through physical attachment. ISME J 11, 641–650 (2017).

5. Durham, B. P. et al. Sulfonate-based networks between eukaryotic phytoplankton and heterotrophic bacteria in the surface ocean. Nat Microbiol 4, 1706–1715 (2019).

6. Christie-Oleza, J. A., Sousoni, D., Lloyd, M., Armengaud, J. & Scanlan, D. J. Nutrient recycling facilitates long-term stability of marine microbial phototroph-heterotroph interactions. Nat Microbiol 2, (2017).

7. Barak-Gavish, N. et al. Bacterial virulence against an oceanic bloom-forming phytoplankter is mediated by algal DMSP. Sci Adv 4, (2018).

8. Segev, E. et al. Dynamic metabolic exchange governs a marine algal-bacterial interaction. Elife 5, (2016).

9. Grossart, H. P., Czub, G. & Simon, M. Algae-bacteria interactions and their effects on aggregation and organic matter flux in the sea. Environ Microbiol 8, 1074–1084 (2006).

10. Amin, S. A. et al. Interaction and signalling between a cosmopolitan phytoplankton and associated bacteria. Nature 522, 98–101 (2015).

11. Mühlenbruch, M., Grossart, H., Eigemann, F. & Voss, M. Mini-review: Phytoplankton-derived polysaccharides in the marine environment and their interactions with heterotrophic bacteria. Environ Microbiol 20, 2671–2685 (2018).

12. Bertrand, E. M. et al. Phytoplankton–bacterial interactions mediate micronutrient colimitation at the coastal Antarctic sea ice edge. Proceedings of the National Academy of Sciences 112, 9938–9943 (2015).

13. Czaran, T. L., Hoekstra, R. F. & Pagie, L. Chemical warfare between microbes promotes biodiversity. Proceedings of the National Academy of Sciences 99, 786–790 (2002).

14. Bidle, K. D. & Falkowski, P. G. Cell death in planktonic, photosynthetic microorganisms. Nat Rev Microbiol 2, 643–655 (2004).

15. Pernthaler, J. Predation on prokaryotes in the water column and its ecological implications. Nat Rev Microbiol 3, 537–546 (2005).

16. Lima-Mendez, G. et al. Determinants of community structure in the global plankton interactome. Science (1979) 348, (2015).

17. Seymour, J. R., Amin, S. A.Raina, J.-B. & Stocker, R. Zooming in on the phycosphere: the ecological interface for phytoplankton–bacteria relationships. Nat Microbiol 2, 17065 (2017).

18. Malfatti, F., Samo, T. J. & Azam, F. High-resolution imaging of pelagic bacteria by Atomic Force Microscopy and implications for carbon cycling. Isme J 4, 427–439 (2010).

19. Bratbak, G. & Thingstad, T. F. Phytoplankton-Bacteria Interactions - an Apparent Paradox - Analysis of a Model System with Both Competition and Commensalism. Marine Ecology-Progress Series 25, 23–30 (1985).

20. Hibbing, M. E., Fuqua, C., Parsek, M. R. & Peterson, S. B. Bacterial competition: surviving and thriving in the microbial jungle. Nat Rev Microbiol 8, 15–25 (2009).

21. Moore, C. M. et al. Processes and patterns of oceanic nutrient limitation. Nat Geosci 6, 701–710 (2013).

22. Thingstad, T. F. et al. Nature of Phosphorus Limitation in the Ultraoligotrophic Eastern Mediterranean. Science (1979) 309, 1068–1071 (2005).

23. Moran, M. A. & Durham, B. P. Sulfur metabolites in the pelagic ocean. Nat Rev Microbiol 17, 665–678 (2019).

24. Thornton, D. C. O. Dissolved organic matter (DOM) release by phytoplankton in the contemporary and future ocean. Eur J Phycol 49, 20–46 (2014).

25. Moran, M. A. et al. Deciphering ocean carbon in a changing world. Proceedings of the National Academy of Sciences 113, 3143–3151 (2016).

26. Zhang, C. et al. Evolving paradigms in biological carbon cycling in the ocean. Natl Sci Rev 5, 481–499 (2018).

27. Horňák, K., Kasalický, V., Šimek, K. & Grossart, H.-P. Strain-specific consumption and transformation of alga-derived dissolved organic matter by members of the Limnohabitans-C and Polynucleobacter-B clusters of Betaproteobacteria. Environ Microbiol 19, 4519–4535 (2017).

28. Kujawinski, E. B. The Impact of Microbial Metabolism on Marine Dissolved Organic Matter. Ann Rev Mar Sci 3, 567–599 (2011).

29. Lechtenfeld, O. J., Hertkorn, N., Shen, Y., Witt, M. & Benner, R. Marine sequestration of carbon in bacterial metabolites. Nat Commun 6, 6711 (2015).

30. Ma, L., Calfee, B. C., Morris, J. J., Johnson, Z. I. & Zinser, E. R. Degradation of hydrogen peroxide at the ocean’s surface: the influence of the microbial community on the realized thermal niche of Prochlorococcus. ISME J 12, 473 (2017).

31. Morris, J. J., Johnson, Z. I., Szul, M. J., Keller, M. & Zinser, E. R. Dependence of the cyanobacterium Prochlorococcus on hydrogen peroxide scavenging microbes for growth at the ocean’s surface. PLoS One 6, e16805 (2011).

32. Morris, J. J., Lenski, R. E. & Zinser, E. R. The black queen hypothesis: Evolution of dependencies through adaptive gene loss. mBio 3, (2012).

33. Paz-Yepes, J., Brahamsha, B. & Palenik, B. Role of a microcin-C-like biosynthetic gene cluster in allelopathic interactions in marine Synechococcus. Proc Natl Acad Sci U S A 110, 12030–12035 (2013).

34. Manage, P. M., Kawabata, Z. & Nakano, S. Algicidal effect of the bacterium Alcaligenes denitrificans on Microcystis spp. Aquatic Microbial Ecology 22, 111–117 (2000).

35. Mayali, X. & Azam, F. Algicidal bacteria in the sea and their impact on algal blooms. Journal of Eukaryotic Microbiology 51, 139–144 (2004).

36. Seyedsayamdost, M. R., Case, R. J., Kolter, R. & Clardy, J. The Jekyll-and-Hyde chemistry of Phaeobacter gallaeciensis. Nat Chem 3, 331–335 (2011).

37. Partensky, F. & Garczarek, L. Prochlorococcus: Advantages and Limits of Minimalism. Ann Rev Mar Sci 2, 305–331 (2010).

38. Scanlan, D. J. et al. Ecological Genomics of Marine Picocyanobacteria. Microbiology and Molecular Biology Reviews 73, 249–299 (2009).

39. Biller, S. J., Berube, P. M., Lindell, D. & Chisholm, S. W. Prochlorococcus: the structure and function of collective diversity. Nat Rev Microbiol 13, 13–27 (2015).

40. Aharonovich, D. & Sher, D. Transcriptional response of Prochlorococcus to co-culture with a marine Alteromonas: Differences between strains and the involvement of putative infochemicals. ISME Journal 10, 2892–2906 (2016).

41. Cubillos-Ruiz, A., Berta-Thompson, J. W., Becker, J. W., Van Der Donk, W. A. & Chisholm, S. W. Evolutionary radiation of lanthipeptides in marine cyanobacteria. Proc Natl Acad Sci U S A 114, E5424–E5433 (2017).

42. Li, B. et al. Catalytic promiscuity in the biosynthesis of cyclic peptide secondary metabolites in planktonic marine cyanobacteria. Proc Natl Acad Sci U S A 107, 10430– 10435 (2010).

43. Biller, S. J., Coe, A. & Chisholm, S. W. Torn apart and reunited: Impact of a heterotroph on the transcriptome of Prochlorococcus. ISME Journal 10, 2831–2843 (2016).

44. Zheng, Q. et al. Dynamics of Heterotrophic Bacterial Assemblages within Synechococcus Cultures. Appl Environ Microbiol 84, (2017).

45. Sher, D., Thompson, J. W., Kashtan, N., Croal, L. & Chisholm, S. W. Response of Prochlorococcus ecotypes to co-culture with diverse marine bacteria. ISME J 5, 1125– 1132 (2011).

46. Cruz, B. N. & Neuer, S. Heterotrophic Bacteria Enhance the Aggregation of the Marine Picocyanobacteria Prochlorococcus and Synechococcus. Front Microbiol 10, (2019).

47. Gärdes, A., Ramaye, Y., Grossart, H. P., Passow, U. & Ullrich, M. S. Effects of Marinobacter adhaerens HP15 on polymer exudation by Thalassiosira weissflogii at different N:P ratios. Mar Ecol Prog Ser 461, 1–14 (2012).

48. Weissberg, O., Aharonovich, D. & Sher, D. Phototroph-heterotroph interactions during growth and long-term starvation across Prochlorococcus and Alteromonas diversity. ISME Journal 17, 227–237 (2023).

49. Seymour, J. R., Ahmed, T., Durham, W. M. & Stocker, R. Chemotactic response of marine bacteria to the extracellular products of Synechococcus and Prochlorococcus. Aquatic Microbial Ecology 59, 161–168 (2010).

50. Roth-Rosenberg, D. et al. Prochlorococcus rely on microbial interactions rather than on chlorotic resting stages to survive long-term stress. mBio 11, e01846–20 (2020).

51. Hennon, G. M. M. et al. The impact of elevated CO2 on Prochlorococcus and microbial interactions with ‘helper’ bacterium Alteromonas. ISME J 12, 520 (2017).

52. Wu, Z. et al. Single-cell measurements and modelling reveal substantial organic carbon acquisition by Prochlorococcus. Nat Microbiol 7, 2068–2077 (2022).

53. Muñoz-Marín, M. C. et al. Mixotrophy in marine picocyanobacteria: use of organic compounds by Prochlorococcus and Synechococcus. ISME J 14, 1065–1073 (2020).

54. del Carmen Muñoz-Marín, M., López-Lozano, A., Moreno-Cabezuelo, J.Á., Díez, J. & García-Fernández, J. M. Mixotrophy in cyanobacteria. Curr Opin Microbiol 78, 102432 (2024).

55. Dubinsky, Z. & Berman-Frank, I. Uncoupling primary production from population growth in photosynthesizing organisms in aquatic ecosystems. Aquat Sci 63, 4–17 (2001).

56. Arnosti, C. Microbial Extracellular Enzymes and the Marine Carbon Cycle. Ann Rev Mar Sci 3, 401–425 (2011).

57. Zubkov, M. V, Tarran, G. A. & Fuchs, B. M. Depth related amino acid uptake by Prochlorococcus cyanobacteria in the Southern Atlantic tropical gyre. FEMS Microbiol Ecol 50, 153–161 (2004).

58. Wu, Z. et al. Modeling Photosynthesis and Exudation in Subtropical Oceans. Global Biogeochem Cycles 35, e2021GB006941 (2021).

59. Cai, R. & Jiao, N. Recalcitrant dissolved organic matter and its major production and removal processes in the ocean. Deep Sea Research Part I: Oceanographic Research Papers 191, 103922 (2023).

60. Grossowicz, M. et al. Prochlorococcus in the lab and in silico: The importance of representing exudation. Limnol Oceanogr 62, 818–835 (2017).

61. Follows, M. J. & Dutkiewicz, S. Modeling Diverse Communities of Marine Microbes. Annual Review of Marine Science, Vol 3 3, 427–451 (2011).

62. Tilman, D. & Kilham, S. S. Phosphate and silicate growth and uptake kinetics of the diatoms Asterionella formosa and Cyclotella meneghiniana in batch and semicontinuous culture 1. J Phycol 12, 375–383 (1976).

63. Stewart, F. M. & Levin, B. R. Partitioning of resources and the outcome of interspecific competition: a model and some general considerations. Am Nat 107, 171–198 (1973).

64. Bar-Yosef, Y., Sukenik, A., Hadas, O., Viner-Mozzini, Y. & Kaplan, A. Enslavement in the water body by toxic aphanizomenon ovalisporum, inducing alkaline phosphatase in phytoplanktons. Current Biology 20, 1557–1561 (2010).

65. Hansell, D. A., Carlson, C. A. & Schlitzer, R. Net removal of major marine dissolved organic carbon fractions in the subsurface ocean. Global Biogeochem Cycles 26, (2012).

66. Carlson, C. A. et al. Dissolved organic carbon export and subsequent remineralization in the mesopelagic and bathypelagic realms of the North Atlantic basin. Deep Sea Research Part II: Topical Studies in Oceanography 57, 1433–1445 (2010).

67. Roshan, S. & DeVries, T. Efficient dissolved organic carbon production and export in the oligotrophic ocean. Nat Commun 8, 2036 (2017).

68. Sher, D., Segrè, D. & Follows, M. J. Quantitative principles of microbial metabolism shared across scales. Nat Microbiol 9, 1940–1953 (2024).

69. Coe, A. et al. Survival of Prochlorococcus in extended darkness. Limnol Oceanogr 61, 1375–1388 (2016).

70. Roth-Rosenberg, D., Aharonovich, D., Omta, A. W., Follows, M. J. & Sher, D. Dynamic macromolecular composition and high exudation rates in Prochlorococcus. Limnol Oceanogr 66, 1759–1773 (2021).

71. Moore, L. R. et al. Culturing the marine cyanobacterium Prochlorococcus. Limnol Oceanogr Methods 5, 353–362 (2007).

72. Morris, J. J., Kirkegaard, R., Szul, M. J., Johnson, Z. I. & Zinser, E. R. Facilitation of robust growth of Prochlorococcus colonies and dilute liquid cultures by ‘helper’ heterotrophic bacteria. Appl Environ Microbiol 74, 4530–4534 (2008).

73. Williams, R. G. & Follows, M. J. Ocean Dynamics and the Carbon Cycle: Principles and Mechanisms. (Cambridge University Press, 2011).

