## Supplementary Information for "Four Potential Mechanisms Underlying Phytoplankton-Bacteria Interactions Assessed Using Experiments and Models"

3) Current address: College of Environmental Sciences and Engineering, State Environmental Protection Key Laboratory of All Material Fluxes in River Ecosystems, Peking University, Beijing, China

### Supplementary Information

#### Supplementary Text 1: Detailed description of the heterotrophs selected for co-culture.

We chose eight heterotrophic bacteria to pair with *Prochlorococcus* MED4 in co-culture, aiming to cover a wide range of previously-describe interactions, from synergistic to antagonistic. Previously, most of the studies of *Prochlorococcus*-heterotroph interactions focused on several strains of the marine gamma proteobacterium *Alteromonas* [1–9]. *Prochlorococcus* has also been studied in co-culture with additional copiotrophic bacteria (*Thalassospira*, *Roseobacter*, *Alcanovorax*, and *Marinobacter*), the oligotroph SAR11 (*Pelagibacter*), and *Synechococcus* [10–12]. In most cases the interaction is mutually beneficial, although *Alteromonas* also inhibits one *Prochlorococcus* strain (MIT9313) in a dose dependent manner, resulting in an extended lag phase [1]. The heterotrophic strains we selected here were aimed to increase the breadth of analyzed *Prochlorococcus*-heterotroph interactions and to focus on strains previously studied in co-culture with other phytoplankton:

*Pseudoalteromonas haloplanktis* is attracted to *Prochlorococcus* exudates, it senses their presence and moves towards them, indicating possible ecological associations [13].

The positive interaction between the Roseobacter *Ruegeria pomeroyi* and the cyanobacteria *Synechococccus*, has been studied extensively [14–16]. An important interaction mechanism is through nutrient exchange. *Synechococcus* produces nitrogen-rich DOM which is consumed by *Ruegeria* and remineralised, producing ammonium, which is quickly used by the *Synechococcus*. *Synechococcus* is sensitive to accumulation of its own DOM and needs *Ruegeria* help to degrade it [15].

Another Roseobacter is *Phaeobacter gallaeciensis* which grows and dies in correlation with blooms of the coccolithophore *Emiliania huxleyi* [17]. Seyedsayamdost at el. showed that *P. gallaeciensis* alternates between mutualistic and antagonistic interactions, switching between promoting algal growth by producing growth stimulants (phenylacetic acid) and protective antibiotics (tropodithietic acid) and killing the algae by producing algaecides [17].

*Sulfitobacter pseudonitzschiae* SMR1 is associated with phytoplankton, as it was isolated from a culture of marine diatom[18]. A related *S. pseudonitzschiae*, SA11, supports the growth of the diatom *Pseudonitzschia multiseries* through the exchange of ammonium and the production of a plant hormone, indole-3-acetic acid (IAA) [19].

Another diatom-associated strain is *Marinobacter adhaerens* HP15, which promotes diatom production of transparent exopolymer particles (TEP) and aggregation [20].

Finally, *Marinovum* HOT5F3 and *Roseovarius* HOT5C3, were studied in the context of co-cultures with *Prochlorococcus* MED4 and MIT9313. In [12] *Marinovum* HOT5F3 inhibited the growth of both *Prochlorococcus* strains, and *Roseovarius* HOT5C3 enhanced the growth of MIT9313 and did not affect that of MED4. The genomes of these two strains have been sequenced and are currently being analyzed.

#### Supplementary Text 2: Model description

**Overall model description**: The four models described in Supplementary Figure S1 were developed to test different hypotheses regarding the mechanisms underlying microbial interactions. Across all models, carbon is introduced into the food web through photosynthesis by autotrophs, and the heterotrophs rely on this organic carbon (no heterotrophic C fixation). The system is closed for N (i.e. no nitrogen fixation), and other nutrients like phosphate and trace metals are not included in the model as they are available in abundance in our experimental media and are unlikely to have significant influence on the growth pattern.

The modeled system is closed for nitrogen, since to the best of our knowledge neither organism can fix dinitrogen or denitrify. Thus, the total amount of nitrogen in the nutrient pools and in biomass is fixed to the initial nitrogen available at the beginning of the experiment. Both get their nitrogen needs from inorganic nitrogen (DIN) such as ammonium. In the basic model formulation, the heterotroph can also uptake organic nitrogen (DON), e.g., amino acids, and nucleotides (which the autotrophs can also utilize in the mixotrophy model). Part of the organic N and C pools are in recalcitrant form (RDOC and RDON), i.e. cannot be utilized and degraded by microorganisms, representing the extremely large pools of refractory DOC and DON in the oceans. Unlike the N pool, the C pool is open, because $CO_2$ dissolves in water or can be released into the air as a function of its concentration in both media. The DIC pool is also replenished when both organisms respire, releasing $CO_2$ (DIC).

Within the cells, N and C are taken up into cellular stores, which are then combined into biomass through biosynthesis. Respiration is from the C store (which is conceptually considered to be formed of e.g. simple carbohydrates), and biomass can be degraded back into C and N stores if C is required for respiration. Organic matter (both labile and recalcitrant) is released from the cells through mortality (in the basic model there is no explicit representation of other processes such as exudation, which is represented only in the overflow model). Finally, DON is slowly spontaneously broken down into DIN, representing an abiotic process of degradation [21].

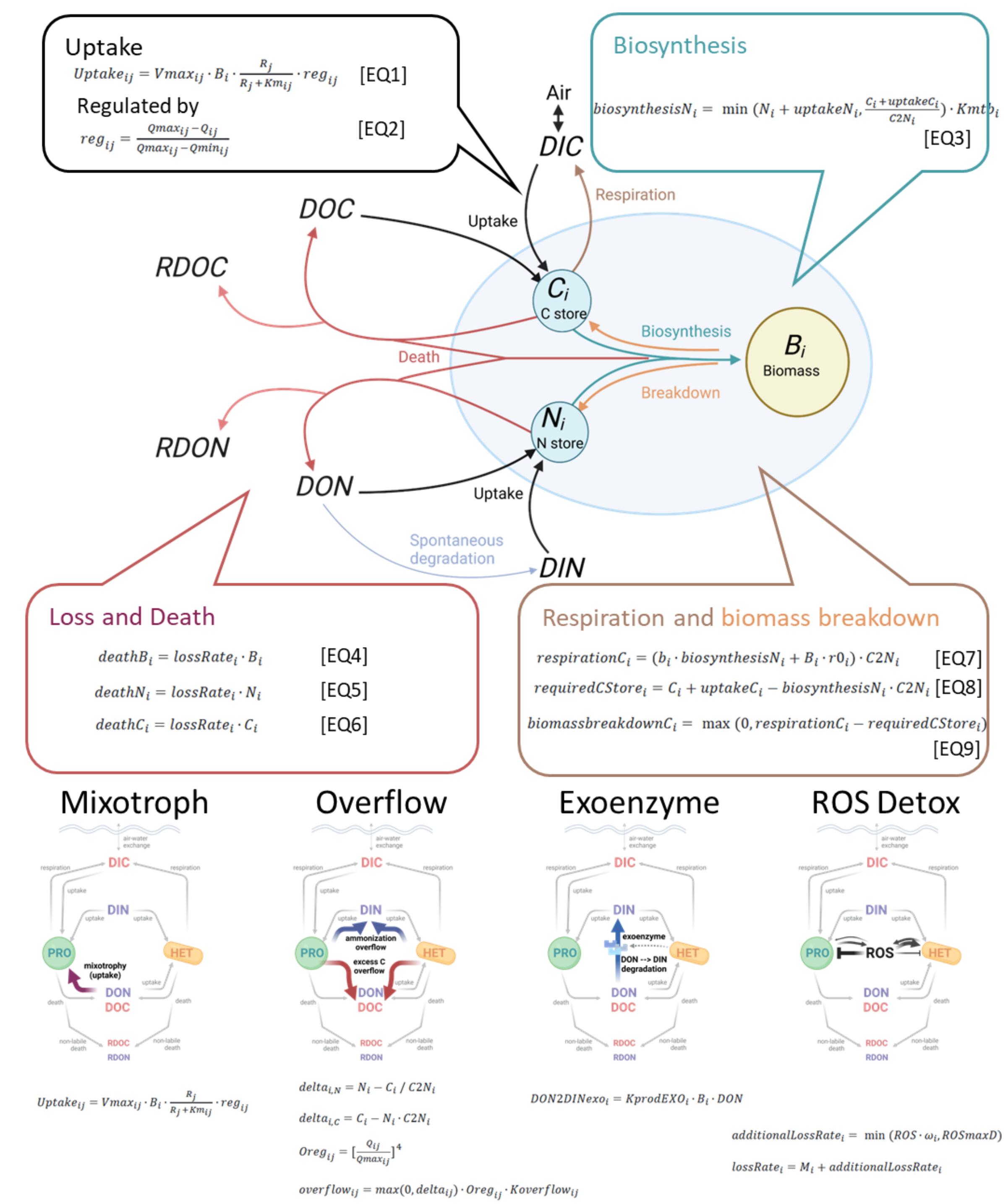


Supplementary Figure S1: Schematic structure of the model, including main model equations and definition of four metabolic mechanisms.

| Variable | Unit | Description |
|---|---|---|
| ***Bi*** | μ M N | Functional biomass in N |
| ***Ni*** | μ M N | N store |
| ***Ci*** | μ M C | C store |
| ***DON*** | μ M N | Dissolved Organic N |

| | | |
|---|---|---|
| ***RDON*** | μ M N | Recalcitrant Dissolved Organic N |
| ***DIN*** | μ M N | Dissolved Inorganic N |
| ***DOC*** | μ M C | Dissolved Organic C |
| ***RDOC*** | μ M C | Recalcitrant Dissolved Organic C |
| ***DIC*** | μ M C | Dissolved Inorganic C |
| ***ROS*** | μ M ROS | Reactive Oxygen Species |

Supplementary Table S1: Model variables

Subscript *i* stands for population, *p* for vPro and *h* for vHet

| **Model** | **Tune able** | **Parameter** | **Units** | **Description** | **Default value** | |
|---|---|---|---|---|---|---|
| | | | | | **PRO** | **HET** |
| **All models** | Yes | $V_{max}DIC_i$ | $s^{-1}$ | C specific inorganic C maximum uptake rate | $5.56*10^{-6}$ | |
| **All models** | Yes | $V_{max}DIN_i$ | $s^{-1}$ | N specific inorganic N maximum uptake rate | $5.56*10^{-6}$ | $6.67*10^{-5}$ |
| **All models** | Yes | $V_{max}DOC_i$ | $s^{-1}$ | C specific organic C maximum uptake rate | $1.85*10^{-6}$ ◊ | $6.67*10^{-5}$ |
| **All models** | Yes | $V_{max}DON_i$ | $s^{-1}$ | N specific organic N maximum uptake rate | $1.11*10^{-6}$ ◊ | $6.67*10^{-5}$ |
| **All models** | Yes | $K_mDIC_i$ | uM C | inorganic C Km (half saturation) | 37 | |
| **All models** | Yes | $K_mDIN_i$ | uM N | inorganic N Km (half saturation) | 0.02 | 0.02 |
| **All models** | Yes | $K_mDOC_i$ | uM C | organic C Km (half saturation) | 0.03 ◊ | 0.1 |
| **All models** | Yes | $K_mDON_i$ | uM N | organic N Km (half saturation) | 0.1 ◊ | 0.1 |
| **All models** | Yes | $M_i$ | $s^{-1}$ | death rate | $1.16*10^{-6}$ | $1.16*10^{-6}$ |
| **All models** | Yes | $\gamma_i$ | no units | % of labile dead matter | 0.8 | 0.8 |
| **All models** | No | KdecayDON | $s^{-1}$ | DON2DIN intrinsic breakdown rate | $2.31*10^{-8}$ | |
| **All models** | No | $Kmtb_i$ | $s^{-1}$ | Maximum metabolic rate | $3.5*10^{-5}$ | $1.8*10^{-4}$ |
| **All models** | No | $QCmax_i$ | ratio | Maximum C:N ratio | 10 | 10 |
| **All models** | No | $QCmin_i$ | ratio | Minimum C:N ratio | 4 | 4 |
| **All models** | No | $R_i$ | ratio | C:N ratio of functional biomass | 6.625 | 5 |
| **All models** | No | $b_i$ | no units | respiration coefficient | 0.01 | 0.01 |
| **All models** | No | $r0_i$ | $s^{-1}$ | dark respiration | $2.08*10^{-6}$ | $2.08*10^{-6}$ |
| **OVERFLOW** | Yes | $Koverflow_i$ | $s^{-1}$ | organic leakage rate | $1.16*10^{-6}$ | $1.16*10^{-6}$ |
| **EXOENZYME** | Yes | $KprodEXO_i$ | $s^{-1}$ uM $N^{-1}$ | DON2DIN Exoenzyme breakdown rate | | $1.16*10^{-6}$ |
| **ROS** | Yes | $KlossROS_i$ | $s^{-1}$ uM $N^{-1}$ | max ROS breakdown | $1.96*10^{-6}$ | $9.72*10^{-6}$ |
| **ROS** | Yes | $KprodROS_i$ | uM H2O2 $s^{-1}$ uM $N^{-1}$ | ROS release rate | $4.7*10^{-9}$ | $1.94*10^{-8}$ |
| **ROS** | Yes | $\omega_i$ | $s^{-1}$ uM $H2O2^{-1}$ | ROS toxicity (additional death) | $3.41*10^{-5}$ | $3.41*10^{-8}$ |

| | | | | | |
|---|---|---|---|---|---|
| **ROS** | No | KdecayROS | $s^{-1}$ | ROS intrinsic decay rate | $2.78*10^{-6}$ |
| **ROS** | No | ROSmaxD | $s^{-1}$ | max ROS toxicity (additional death) | $3.21*10^{-5}$ |

Supplementary Table S2: Model parameters

◊ - *Prochlorococcus* Organic growth parameters, used in Mixotroph model only.

#### General model description

Phytoplankton and bacteria are modeled using the approach of [22], with 3 variables: N store ($N_i$), C store ($C_i$) and biomass ($B_i$) (Supplementary Figure S1), with the subscript *i* representing each of the interacting organisms. C and N are taken up from the corresponding nutrient pools into the C and N stores respectively, which are then combined through biosynthesis into functional biomass (equation 1). This model formulation supports variable C:N ratios as the N and C stores are maintained independently. The model equations are formalized in units of nitrogen, with the C:N ratio $C2N_i$ used every time a process or pool involving C needs to be translated into nitrogen units.

The functional biomass has a fixed C:N ratio ($C2N_i$). Biomass can be synthesized, broken down to provide C for respiration, or released due to death:

$$\mathrm{dB_i/dt = biosynthesisN_i - biomassbreakdownC_i/C2N_i - deathB_i} \quad (1)$$

The stores are utilized for biosynthesis, respiration (C store only) and are degraded into dissolved organic matter upon death (equations 2, 3). When the stores are imbalanced, some of them may be exuded as overflow (meaning that *overflowCi* in equation 3 is set to zero in all models apart from the overflow one, see below).

$$dN_i/dt = uptakeN_i + biomassbreakdownC_i/C2N_i - biosynthesisN_i - overflowN_i - deathN_i \quad (2)$$

$$dC_i/dt = uptakeC_i + biomassbreakdownC_i - biosynthesisN_i \cdot C2N_i - respirationC_i - overflowC_i - deathC_i \quad (3)$$

#### Carbon Resources

Carbon resources are split into inorganic (DIC), organic (DOC), and recalcitrant, non-labile organic (RDOC). The carbon system is not closed, as new DIC is absorbed and lost from the air via air-water exchange of $CO_2$.

$$dDIC/dt = DICAirWaterExchange + \sum_{i\in\{p,h\}} \left(respirationC_i - grossUptake_{i,DIC}\right) \quad (4)$$

$$dDOC/dt = \sum_{i\in\{p,h\}} \left(totaldeathC_i \cdot \gamma_i + overflowC_i - grossUptake_{i,DOC}\right) \tag{5}$$

$$dRDOC/dt = \sum_{i\in\{p,h\}} totaldeathC_i \cdot (1-\gamma_i) \tag{6}$$

$$totaldeathC_i = deathB_i \cdot C2N_i + deathC_i \tag{7}$$

Here $\gamma_i$ is the portion of the losses (death and leakiness) that is labile (avaliable for consumption).

DIC can also be exchanged with air, replenishing the C supply

$$DICAirWaterExchange = -\frac{\dfrac{DIC - csat}{h}}{Kg \cdot B \cdot 0.01} \tag{8}$$

Here $csat$ is the saturated DIC concentration[23], $h$ is the height of the media in meters, $Kg$ is the exchange rate in m sec$^{-1}$ (measured in the lab, see methods), and $B$ is the Revelle buffer factor [24].

#### Nitrogen Resources

The nitrogen budget consists of the initial nitrogen in organic (DON), inorganic (DIN) and recalcitrant, non-labile organic (RDON), as well as the nitrogen in the cell biomass and nitrogen store. Thus, the system as a whole is closed for nitrogen. DON slowly decays abiotically into DIN (in all models). In the EXOENZYME model it is also degraded by exoenzymes released by the heterotroph (see DON breakdown due to exoenzymes)

$$dDIN/dt = \sum_{i\in\{p,h\}} \left(overflowN_i + DON2DIN_i - grossUptake_{i,DIN}\right) + globalDON2DIN \tag{9}$$

$$dDON/dt = \sum_{i\in\{p,h\}} \left(totaldeathN_i \cdot \gamma_i - grossUptake_{i,DON} - DON2DINexo_i\right) - globalDON2DIN \tag{10}$$

$$dRDON/dt = \sum totaldeathN_i \cdot (1-\gamma_i) \tag{11}$$

$$totaldeathN_i = deathB_i + deathN_i \tag{12}$$

$$globalDON2DIN = KdecayDON \cdot DON \tag{13}$$

Here, $KdecayDON$ is the rate of DIN abiotic decay into DON.

#### Nutrient uptake

Nutrient uptake is modeled as follows, using Monod/Michaelis-Menten kinetics regulated by the internal stores of each element:

$$grossUptake_{ij} = Vmax_{ij} \cdot B_i \cdot \frac{R_j}{R_j + Km_{ij}} \cdot reg_{ij} \tag{14}$$

Where $R_j$ is one of the resources ($DOC, DON, DIC, DIN$), $Vmax_{ij}$ stands for the maximum uptake rate, $Km_{ij}$ is the half-saturation constant and $reg_{ij}$ quantifies the regulation of nutrient uptake, enabling the model to capture variable C:N ratios while avoiding wildly imbalanced N and C stores:

$$reg_{ij} = \frac{Qmax_{ij} - Q_{ij}}{Qmax_{ij} - Qmin_{ij}} \tag{15}$$

In equation 15, for a given strain *i* and a given nutrient *j*, $Qmax_{ij}$ and $Qmin_{ij}$ are the minimum and maximum nutrient ratios for this organism, and $Q_{ij}$ is the current nutrient ratio. $reg_{ij}$ is also clipped to be between 0 and 1. This function approaches 0 as the ratio reaches the max value, curtailing uptake.

$$Q_{i,C} = \frac{C_i + B_i \cdot C2N_i}{N_i + B_i} \tag{16}$$

$$Q_{i,N} = \frac{N_i + B_i}{C_i + B_i \cdot C2N_i} \tag{17}$$

The heterotroph has 2 sources of nitrogen, DIN and DON. In theory, when it is carbon limited, it can prioritize the uptake of one of these nitrogen pools over the other, e.g. if there is a clear energetic advantage to utilizing organic vs inorganic forms. However, since we have no empirical data on this for *Prochlorococcus* or marine heterotrophic bacteria, in our model the heterotroph will uptake as much as possible from each pool (and the autotroph will do the same in the specific case where mixotrophy is modelled).

#### Biosynthesis and respiration

Biosynthesis is the process of creating functional biomass (e.g. proteins, nucleic acids, lipids etc.) out of the stores. Biosynthesis is constrained by the size of the smaller of the stores, through equation 18:

$$biosynthesisN_i = \min\left(N_i + uptakeN_i, \frac{C_i + uptakeC_i}{C2N_i}\right) \cdot Kmtb_i \tag{18}$$

Here, $Kmtb_i$ represents the biosynthesis rate. The functional biomass of our organism has a fixed C:N ratio, determined by values in the literature (C:N ratio of 5 for heterotrophs and Redfield ratio of 6.625 for the autotrophs, [25]). Together with the variable C and N stores this allows a variable overall elemental ratio.

All living organisms respire, releasing DIC. We model both maintenance and growth-related respiration [26]. Carbon for respiration is initially drawn from the C store (which can be thought of as carbohydrates or glycogen), and if the C store is depleted, the cells break down sufficient biomass for respiration needs [22].

$$respirationC_i = (b_i \cdot biosynthesisN_i + B_i \cdot r0_i) \cdot C2N_i \tag{19}$$

Biomass may be broken down to support respiration. Respiration is assumed to occur from C storage (e.g. carbohydrates or lipids), but if there is not enough C storage biomass will be degraded instead (equation 20,21). Note that equation 20 is calculated after the calculation of biosynthesis but before the calculation of the C store, and hence the needs to incorporate also the C used for biosynthesis.

$$requiredCStore_i = C_i + uptakeC_i - biosynthesisN_i \cdot C2N_i \tag{20}$$

$$biomassbreakdownC_i = \max(0, respirationC_i - requiredCStore_i) \tag{21}$$

#### Death and the effect of ROS

We model death as an exponential process, where a fixed fraction of the population dies every day. In a previous study [2] we show that other mathematical formulations of mortality are better at describing the decline of *Prochlorococcus-Alteromonas* cocultures, but these are emergent properties of the co-culture which we are explicitly testing here, and for the axenic cultures the exponential decay provided a good fit. Most of the biomass released during death is in the form of organic matter (DOC and DON), however, a portion of this organic matter is recalcitrant, i.e. cannot be recycled or re-used by the organisms in co-culture (RDOC and RDON). The recalcitrant pool grows with time, reducing the bioavailable N in the system (DIC enters the systems through water-air exchange and is fixed from photosynthesis, hence the C pool is open). Death also implicitly models other loss processes, e.g., passive leakage and diffusion, which are explicitly considered separately in the overflow model.

$$deathB_i = lossRate_i \cdot B_i \tag{22}$$

$$deathN_i = lossRate_i \cdot N_i \tag{23}$$

$$deathC_i = lossRate_i \cdot C_i \tag{24}$$

In the model explicitly representing the dynamics of Reactive Oxygen Species (ROS, see details below), ROS is a toxin, killing some of the bacteria, and ROS influence is modeled as additional mortality. The additional mortality is capped by $ROSmaxD$, the saturated maximum toxicity of ROS (see additional details below).

$$additionalLossRate_i = \min(ROS \cdot \omega_i, ROSmaxD) \tag{25}$$

$$lossRate_i = M_i + additionalLossRate_i \quad (26)$$

#### Mechanisms represented in specific models

Four different mechanisms of interaction have been implemented in four separate model variants. Each of these mechanisms was tested to see if it can reproduce the lab experiments.

**Mixotrophy**: In many classic oceanographic models phototrophs are assumed to utilize only inorganic carbon and nitrogen, with the organic forms recycles and remineralized by heterotrophic bacteria [27]. However, *Prochlorococcus* can utilize a wide range of organic molecules [28]. and requires mixotrophy to survive extended periods of dark in axenic cultures[8,29], and at depths where there is insufficient light for photosynthesis in the ocean [22]. In the mixotroph model, *Prochlorococcus* can feed on both inorganic and organic C and N. This is implemented by a non-zero $V_{max}$ for *Prochlorococcus* uptake of DOC and DON in equation 14.

**Overflow**: Overflow metabolism is the process where the uptake of one element in excess of another leads to an elemental imbalance, beyond the capacity of the cell to deal with through increased internal storage pools. Overflow metabolism has been studied in detail in phototrophs, which continue to photosynthesize even when growth is not possible due to a limitation of e.g. N or P, releasing photosynthate (DOC) into the surrounding media [30,31]. Heterotrophic bacteria, under C limitation, may extract energy and carbon from organic compounds such as amino acids, exuding the nitrogen containing waste (e.g. $NH_4^+$, [15,32]). In the overflow model, both organisms release some of the excess stores when their C:N ratio is out of balance (equation 27). Carbon is released to DOC and nitrogen is released as DIN.

$$overflow_{ij} = max\left(0, delta_{ij}\right) \cdot Oreg_{ij} \cdot Koverflow_{ij} \quad (27)$$

Here $delta_{i,N}$ is the excess in the respective N or C store, $Oreg_{ij}$ is a term that regulates the overflow release (see below), and $Koverflow_i$ is the overflow rate ($sec^{-1}$).

$$delta_{i,N} = N_i - C_i/C2N_i \quad (28)$$

$$delta_{i,C} = C_i - N_i \cdot C2N_i \quad (29)$$

Overflow is regulated by a power-of-4 function. This regulation function is sharper, as the organism is only motivated to release overflow under starvation, when it is nearing its max ratio limit, therefore we chose to use power of 4, similar to (Wu et al., 2022).

$$Oreg_{ij} = \left[\frac{Q_{ij}}{Qmax_{ij}}\right]^4 \quad (30)$$

**Exoenzymes**: DON is comprised of a variety of molecules, including high molecular weight compounds which need to be degraded by extracellular enzymes before they can be taken up by bacteria [32]. In our model, we describe this process through the degradation of DON to DIN by exoenzymes released by the heterotroph, with the resulting DIN available for uptake by both organisms. As such, this is a common good service that the heterotroph is providing. In our simplified representation, there is no metabolic cost for the bacteria to release these enzymes, since this cost has to the best of our knowledge not been quantified. We also note that this representation does not truly capture the fact that most molecules released by exoenzymes are not DIN but rather DON (e.g. free amino acids released from proteins during degradation by exoproteases). Representing this process in more molecular detail would require a more highly resolved model. Finally, we represent only DON and not DOC degradation, since the model focuses on the long-term dynamics under N stress.

$$DON2DINexo_i = KprodEXO_i \cdot B_i \cdot DON \tag{31}$$

**ROS detoxification**: In addition to the nutrient exchange mechanisms, another form of potential interaction between phototrophs and heterotrophs is the detoxification of waste products, and specifically ROS. ROS detoxification has been shown to be one of the mechanisms whereby heterotrophic bacteria can support the growth of *Prochlorococcus* and *Synechococcus*[9,33,34]. In the ROS detoxification model, both phytoplankton and bacteria release ROS, break down ROS and die due to ROS toxicity, as described above. The concentration of ROS depends on ROS production rate ($KprodROS_i$), ROS degradation rate ($KlossROS_i$) and ROS intrinsic decay rate ($ROSdecayRate$):

$$dROS/dt = \sum_{i \in \{p,h\}} (ROSrelease_i - ROSbreakdown_i) - ROSdecay \tag{32}$$

$$ROSbreakdown_i = KlossROS_i \cdot B_i \cdot ROS \tag{33}$$

$$ROSrelease_i = KprodROS_i \cdot B_i \tag{34}$$

ROS is intrinsically unstable and decays abiotically

$$ROSdecay = ROSdecayRate \cdot ROS \tag{35}$$

#### Supplementary Text 3: Selection of initial parameter values

We started by estimating the initial parameters for each of the models described above from literature studies. Using these parameters, we were unable to reproduce *Prochlorococcus* growth in axenic cultures (Supplementary Figure S2A). Sensitivity analysis to individual parameters did not identify a single parameter value that, when modified, allowed models to recapitulate axenic growth (Supplementary Figure S2B,C, Supplementary Figure S3), in agreement with previous studies [35]. In this supplementary text section we detail the sources of the initial parameters, with following sections detailing the sensitivity analyses and approach to select the final parameters for the tens of thousands of virtual *Prochlorococcus*-heterotroph pairs.

##### *Estimating $K_m$ and $V_{max}$ for Prochlorococcus and heterotrophic bacteria*

The model employed here uses Monod/Michaelis-Menten equations for the uptake of inorganic and organic C and N. There are few actual measurements of the kinetic parameters $V_{max}$ and $K_m$, and therefore we needed to estimate them from related literature values.

*Prochlorococcus and heterotroph inorganic N*

We used measured uptake rates for ammonium, as this is the most commonly used inorganic N form for *Prochlorococcus*. These rates were taken from Sup Table S2 in Berthelot et al [36] as N-specific $NH_4^+$ uptake rates (0.01 $h^{-1}$). Since the $NH_4^+$ in this study was added as a tracer (30nM), these values represent actual uptake (V). We assume that the $K_m$ for $NH_4^+$ is close to the concentrations found in situ (10-30nM), in agreement with estimations from a model [37]. Since the $K_m$ is close to ambient concentration, the $V_{max}$ should be twice that of V. We note that the uptake rates differ between studies, e.g. the uptake rate in the work of Matsuda and colleagues [37] is 2.42 $d^{-1}$, which is an order of magnitude higher, and higher than the value from the work of Maranon and colleagues is $V_{max}$ $6.7*10^{-5}$ pgN $cell^{-1}$ $h^{-1}$ [38]. The $K_m$ estimated from a genome-scale model are even higher ($K_m$=390uM and $V_{max}$= 0.9 mmol $gDW^{-1}$ $h^{-1}$, [30]). This highlights the difficulty in estimating these parameters.

For the heterotrophs, using the same sources of data and assumptions, N-normalized $NH_4^+$ uptake would have been 0.003 $h^{-1}$[36], which implies a similar $K_m$ and $V_{max}$ per cell as *Prochlorococcus* (given that the N $cell^{-1}$ for heterotrophs is ~3 fold higher for our model). However, these measurements were taken under oceanic conditions probably dominated by slow growing bacteria such as SAR11. The strains used in our experiments replicate at least 4-5 times per day in lab conditions when given ammonium as N source. Hence, we selected $V_{max}$ which is 40 fold higher than in the field, 4-5 $d^{-1}$.

*Prochlorococcus and heterotroph organic N*

We use the uptake rates for leucine in Berthelot et al paper [36]. Here, leucine was added at saturating concentrations hence the "potential V" is $V_{max}$ (0.2-0.5 amol N $cell^{-1}$ $h^{-1}$). This is in the range of concentrations from Table 1 in [39] for free living bacteria: 0.35-0.53

*Prochlorococcus and heterotrophs organic C*

To the best of our knowledge there are no measurements of the growth rate of *Prochlorococcus* on DOC - molecules such as glucose and pyruvate, which are taken up and utilized, do not induce faster growth and cannot serve as sole C sources, [8]. Thus, this parameter cannot be constrained from cultures. We have previously shown that about 80% of the growth of *Prochlorococcus* at depths of ~100m in the oceans is fueled by DOC uptake [22]. Assuming the growth rate of *Prochlorococcus* at these depths is about 0.2 $day^{-1}$, we can assume that the DOC uptake rate needed to fuel this is 0.2*0.8=0.16 $day^{-1}$, which we set as the initial $V_{max}$ for *Prochlorococcus*. For the heterotrophs, similar to the inorganic nitrogen, we set $V_{max}$ to be 4-5 $d^{-1}$ to represent the growth observed in C-replete lab cultures.

We note that the initial estimates of $V_{max}$ we used are all much higher than the actual measurements for glucose in the field, which is often used as a representative organic carbon source. Estimates of $V_{max}$ for glucose range between 0.07-0.12 amol $cell^{-1}$ $h^{-1}$ for heterotrophs (Ayo et al) and 0.4-6 pmol $min^{-1}$ mg $prot^{-1}$ for *Prochlorococcus* [40], assuming 100 fg protein $cell^{-1}$ [41]). These are 4-5 orders of magnitude too low to support observed growth rates in the lab, likely representing the fact that a) glucose is a small fraction of the labile OC; b) These measurements are likely primarily from slow-growing cells in nature, which could also be co-limited by N or P.

For $K_m$ values we chose to use those for glucose from field experiments: 0.03 μM for *Prochlorococcus*, (range 0.015-0.056 μM, [40]), and 0.1 μM for the heterotrophs (range 0.025-2.53μM, [39]).

*Prochlorococcus DIC uptake*

Values were taken from [42] (Table 1, photosynthesis rates). $K_p$=37 $\mu$M, $P_{max}$ = 1.32*$10^{-20}$ mol $cell^{-1}$ $s^{-1}$). Compare with C-specific C fixation rates in Berthelot et al 0.01-0.02 $h^{-1}$ (50-100 amol C cell $h^{-1}$) [36]. Compare with 1.23 fg C $cell^{-1}$ $h^{-1}$ from [43].

*Estimating Exoenzyme Activity*

We model the degradation of DON to DIN as pseudo-first-order kinetics. It is difficult to find appropriate starting values for the rate of exoenzyme activity. $V_{max}$ values from biochemical analyses such as aminopeptidase activity in marine waters from [44] range between 80-700 amol $h^{-1}$ $cell^{-1}$ (~2.5*$10^{-5}$ to 2.2*$10^{-4}$ $sec^{-1}$ $\mu$M $N^{-1}$). These values are 3-4 orders of magnitude higher than rates of $NH_4^+$ turnover rate measured in the field [21] or estimated from the reduction of DON and increase in $NH_4^+$ from [15] (6*$10^{-8}$ to 6*$10^{-6}$ and ~3*$10^{-7}$ $sec^{-1}$ $\mu$M $N^{-1}$, respectively). Additionally, these rates would be 4-5 orders of magnitude higher than the DON uptake rates, assuming a concentration of 10$\mu$M, and thus exoenzymatic activity would dominate the nitrogen dynamics in the model. We therefore selected a value of 1.16*$10^{-6}$ $sec^{-1}$ $\mu$M $N^{-1}$ as a starting point.

*Estimating ROS parameters*

In this study we assume ROS to be $H_2O_2$, as $H_2O_2$ degradation has been studied intensively as one potential mechanism enabling "helper" bacteria to support *Prochlorococcus* survival [9]. While there are other ROS species, including some that have been measured in *Prochlorococcus* and heterotrophic bacteria (e.g. Superoxide, [45]) the much longer half-life of $H_2O_2$ makes it more likely to accumulate in the media and affect the surrounding cells.

Our model has five ROS-related parameters:

1) ROS exudation/production – was parameterized for the heterotrophs from (Figure 4 in [46]) as 1 amol $cell^{-1}$ $hour^{-1}$. There are no published measurements of $H_2O_2$ production for *Prochlorococcus* (but see ROS degradation, below). However, published superoxide production rates (0.003-0.050 amol $cell^{-1}$ $h^{-1}$, [45] are an order of magnitude lower than those of most heterotrophic bacteria (Figure 4 in [46]). Hence, we used 0.1 amol $cell^{-1}$ $h^{-1}$ for *Prochlorococcus* $H_2O_2$ production.
2) ROS degradation – was modelled as a (pseudo)-first order process [46], allowing us to use the values for heterotrophs presented in this publication. Rates for heterotrophs ranged from 2.2*$10^{-8}$-37*$10^{-8}$ $h^{-1}$ [cell $ml^{-1}$]$^{-1}$ (not including Vibrio sp. A535 and Bacillus sp. AzsLept-1c which were clear outliers). We therefore used a value of 10*$10^{-8}$ $h^{-1}$ [cell $ml^{-1}$]$^{-1}$, which is near the median and was measured from *R. pomeroyi* DSS-3, one of the strains we used in our study. ROS degradation for *Prochlorococcus* was measured by [33] and was ~0.10 $d^{-1}$ at 22 degrees C and an initial $H_2O_2$ concentration of 200nM (found in autoclaved Pro99). We assume 100,000 cell $ml^{-1}$ in these experiments, which started at 30,000 cell $ml^{-1}$ and replicate every 1-2 days.
3) The rate of ROS intrinsic degradation is taken from [47].
4) ROS penalty/toxicity – this was estimated from the effect of different concentrations of $H_2O_2$ on the decline rates of *Prochlorococcus* [34]. We first calculated the decline rates from the slope of change in cell numbers over time. We then calculated a value for the effect of $H_2O_2$ on mortality "additional mortality rate" from the regression of mortality rate vs $H_2O_2$ concentration, assuming a growth rate of 0.4 $d^{-1}$ and a "base death" (corresponding to $M_p$ in our model) rate of 0.1 $d^{-1}$. From the same study we took a saturating value of additional mortality at 10mM $H_2O_2$ of 3.2*$10^{-5}$ $sec^{-1}$.

#### Supplementary Text 4: Initial parameterization, sensitivity analysis and generation of virtual *Prochlorococcus* (vPros).

Having selected initial parameter values, we determined to what extent these values can reproduce the growth of axenic *Prochlorococcus* cultures growing in both low-N Pro99 and regular Pro99 (100µM and 800 µM $NH_4^+$, respectively). In the first simulations, *Prochlorococcus* was starved for carbon, and did not grow (Supplementary Figure S2A). Therefore, we doubled *Prochlorococcus* inorganic carbon specific max uptake rate $V_{max}ICp$ to 0.48 $d^{-1}$ which is in the upper range of published data and can theoretically support the growth rate measured in the lab, assuming all carbon is from photosynthesis. With this new value *Prochlorococcus* were able to grow but slowly, reaching max growth on day ~50 instead of day 10 in the lab (Supplementary Figure S2B). We next tested whether changes in a single parameter in addition to the $V_{max}ICp$ can provide good fits to the experimental data. This sensitivity analysis was run by simulating growth while changing each parameter within a range of 2 orders of magnitude (0.1-10X, except for the fraction of recycling of dead matter ($\gamma_i$) where we checked values between 0.1-0.9). No single parameter was sufficient to obtain a good model-data fit (Supplementary Figure S2C, S3).

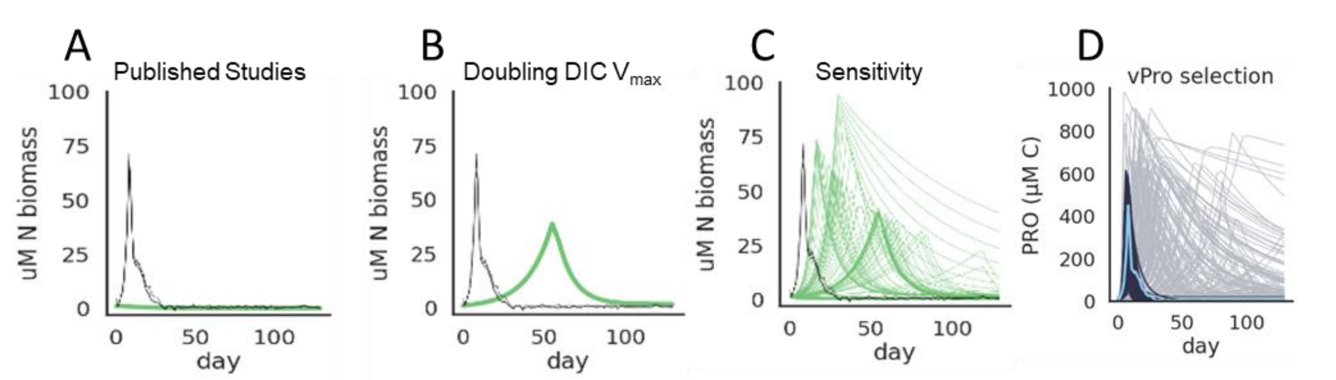


Supplementary Figure S2: Selecting parameters for vPros. A-D) Stages in the process of parameterizing thousands of vPros. A) . no growth using parameters from published studies. B) growth when doubling $V_{max}IC$. C) Sensitivity analysis – no single parameter matches the experimental measurements. D) 0.5% of Montecarlo simulations that match the phenotype

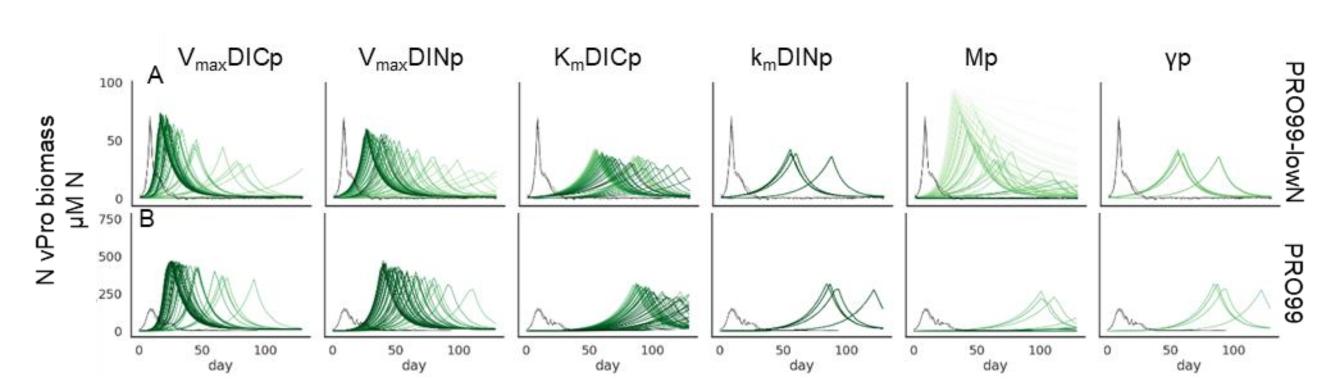


Supplementary Figure S3: Sensitivity of *Prochlorococcus* parameters, changes in *Prochlorococcus* simulation curves when parameter values are changed 2 orders of magnitude (0.1X-10X). *Prochlorococcus* was simulated standalone. A) using pro99-lowN media. B) using Pro99 media.

We next used a Monte-Carlo approach to randomly modify 2 or more parameters from the initial values, generating ~10,000 parameter combinations for each model, and selecting those that best fit the experimental data (Supplementary Figure S2D). Only 0.6% of the simulations (198/49866) matched the lab results for both Pro99 and low-N Pro99, defined as the geometric mean of the Root Mean Square Error (RMSE) on both media being lower than 60. Most of the simulations did not match the lab measurements because *Prochlorococcus* grew poorly, that is, they did not grow (47%), had very weak growth (7%) or growth was too slow and delayed (19%). Another set of simulated *Prochlorococcus* grew too well and achieved unrealistically high biomass (27%). The parameter combination where the simulation results matched the lab for both media were defined as "virtual *Prochlorococcus*" (vPro). This set of vPros is presented in Figure 2.

To obtain more vPros and thus a better representation of the parameter space resulting in specific co-culture outcomes, we performed a second set of random simulations, where we started from a randomly selected vPro from those described above and randomly changed 3-5 of its parameters. This resulted in 300-700 vPros per model, which were used to generate the results presented in Supplementary Figure S12.

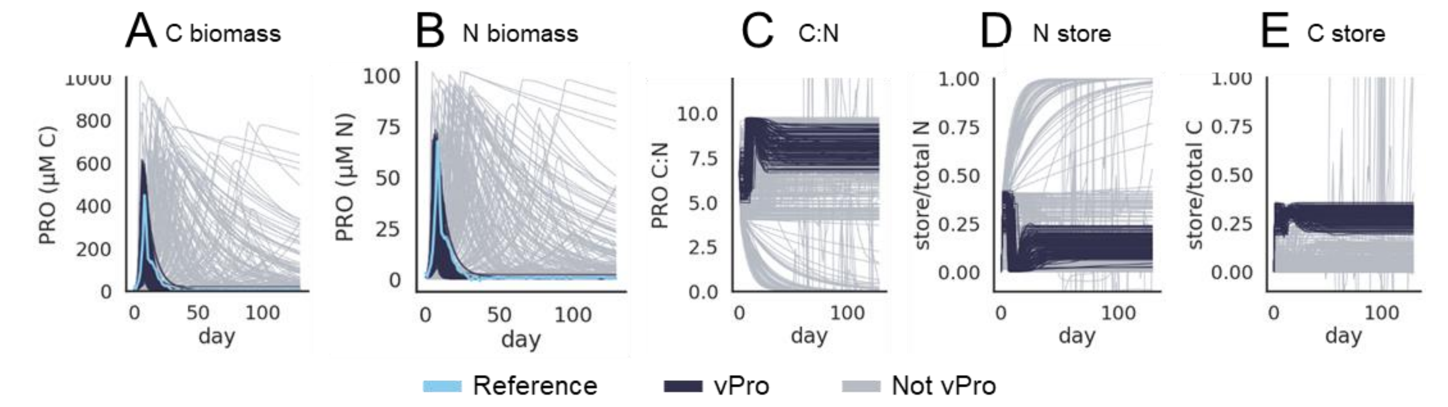


Supplementary Figure S4: vPro Selection based on standalone simulations. A) total vPro C biomass B) total vPro N biomass C) C:N ratio of the simulations D) fraction of the N store from total C biomass. E) fraction of the N store from total N biomass. Black represent selected vPros, light blue – reference experimental measurements, grey – simulations that did not meet vPro criteria

To determine whether the vPros are biologically realistic, we checked whether their predicted C:N ratios are within a realistic range [35]. All selected vPros had selected C:N ratios between 4-10 (mean 7.9±1.3), and the N and C store took up at most 40% of the total biomass (mean 14%±11%, 27%±7% for N and C respectively) (Supplementary Figure S4).

#### Supplementary Text 5: Which parameter combinations result in a good fit to axenic growth curves, defined as vPros?

In the lab, monocultures of *Prochlorococcus* grew exponentially, followed by a sharp decline. This is reflected in the parameter values of the vPros in our model. All vPros from all models have high maximum uptake rates ($V_{max}$) for both DIC and DIN (~2.7 $d^{-1}$), coupled with high mortality (0.5 $d^{-1}$) (Supplementary Figure S5A, S6A). The high values of $V_{max}$ are required to support the exponential growth at the beginning of the simulation. The $V_{max}$ values for inorganic N and C are negative correlated (pearsonr -0.3, pvalue <1e-7), and there seems to be a tradeoff between them. The high death rate is required to reproduce the fast population decline once nutrients are depleted.

One exception is the ROS model, where there is additional mortality, caused by ROS toxicity, and as a result the basal death rate can stay low. In all the models, death rate is correlated with both $V_{max}$es, (pearsonr 0.3-0.4), higher $V_{max}$ is correlated with higher death rate (Supplementary Figure S5, S6).

The selection of vPros is not sensitive to the values of other *Prochlorococcus* parameters, for example, $K_m$ values, parameters related to uptake of organic nutrients, Overflow rates, rates of recycling – all these have low impact on vPro selection in all models (Supplementary Figure S6).

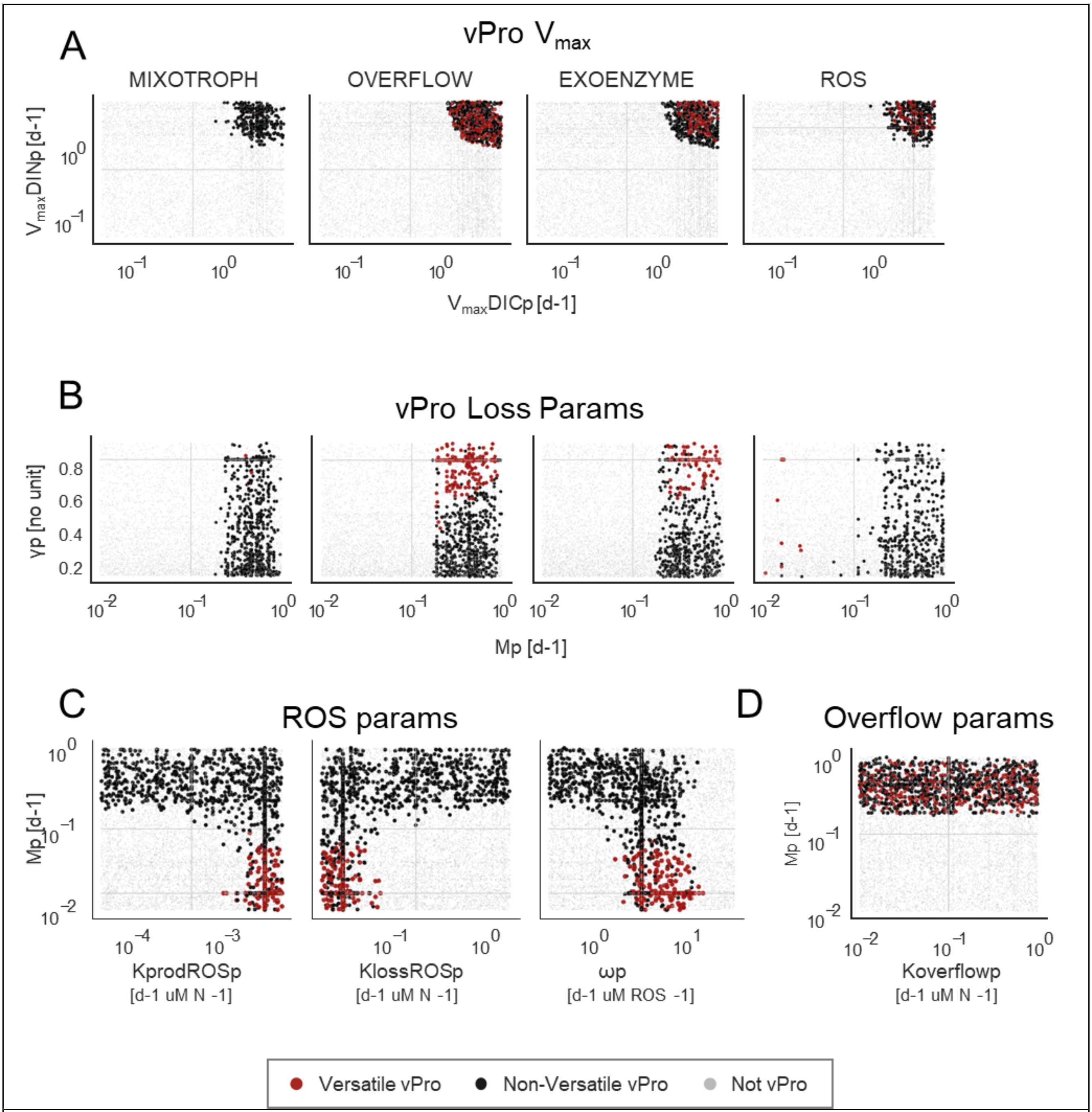


Supplementary Figure S5: Correlation between selected vPro parameters. A) growth parameters. $V_{max}$ICp: vPro max DIC uptake rate, $V_{max}$INp: vPro max DIN uptake rate. B) loss parameters. Mp: vPro intrinsic death rate, $\gamma_p$: fraction of dead matter that is bioavailable. C) ROS parameters. KprodROSp: vPro ROS production rate, KlossROSp: vPro ROS degradation rate, $\omega_p$: vPro susceptibility to ROS toxicity. D) Overflow parameters. Koverflowp: vPro rate of excess overflow release.

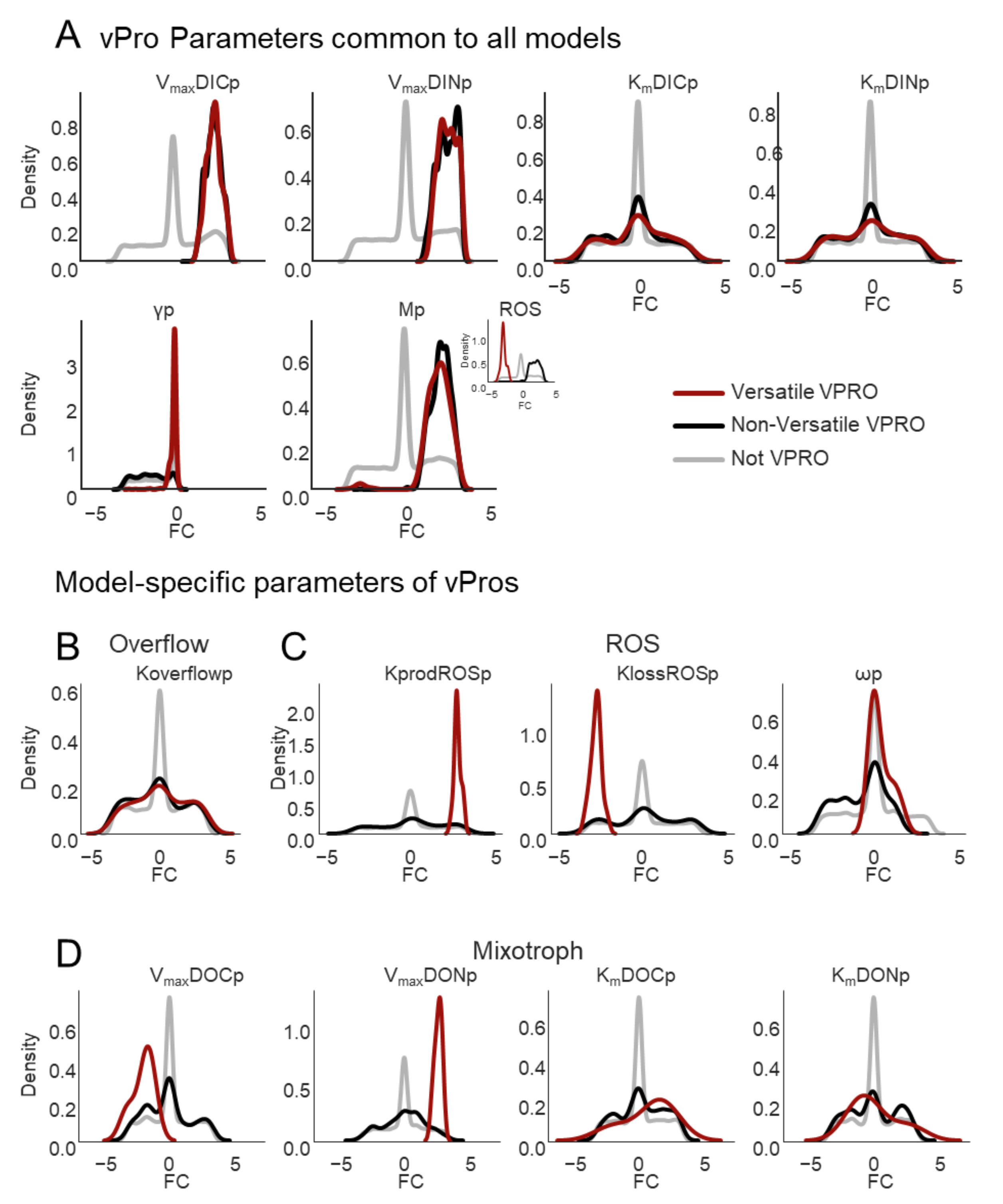


Supplementary Figure S6: Parameters of selected vPros and of versatile vPros. A) vPro parameters common to all models (showing distribution across all models, inset shows ROS death rate). B-D) model-specific vPro parameters B) Overflow vPro parameters. C) ROS vPro parameters. D) Mixotroph vPro parameters. Exoenzyme does not have model specific vPro parameters.

#### Supplementary Text 6: Classification of simulations as specific co-culture outcomes

Having selected several hundred vPros for every model, we next generated ~10,000 virtual heterotrophs (vHets) with parameter combinations randomly selected from the ranges described above, and simulated their growth in pairwise co-cultures with one of the vPros. We then asked whether each of the vPro-vHet simulations is similar to one of the co-culture outcomes, or whether it is different from all of them ("other"). Our approach to this classification problem was to use a stacking classifier, described in detail below, which combines two logistic regression classifiers (separately for N and C biomass) and a random forest classifier based on a set of features describing the growth curves.

As training sets for the classifier, we used the growth curves from this study (Figure 1B) and from a previous study of *Prochlorococcus* mortality in co-culture with different *Alteromonas* strains [2]. We also used another dataset where multiple strains of *Prochlorococcus* were grown axenically using different media, which we used to create 'Other' group, representing curves not covered in our (Soussan-Farhat et al, manuscript in prep). For the training set, we interpolated the data to align the measurement times in all samples, clipped values below 1 µM N or C as this is the experimental limit of detection in our reference data and used only the first 90 days to fit the shortest curves available.

The initial training set contained 417 samples but was small and highly unbalanced (Supplementary Table S3, Supplementary Figure S7). We therefore used SMOTEENN from imbalanced-learn Python package to remove this imbalance by automatically creating additional training samples similar to the experimental measurements and to remove 'Other' samples that were too close to our phenotypes. The final training set contained 1,747 curves.

| Phenotype | number of training samples | number of training samples after fixing imbalance |
|---|---|---|
| **Other** | 333 | 82 |
| **Inhibited** | 36 | 333 |
| **Axenic** | 18 | 333 |
| **Strong** | 18 | 333 |
| **Sustained** | 6 | 333 |
| **Weak** | 6 | 333 |
| **Total** | 417 | 1747 |

Supplementary Table S3: The number of training samples per class

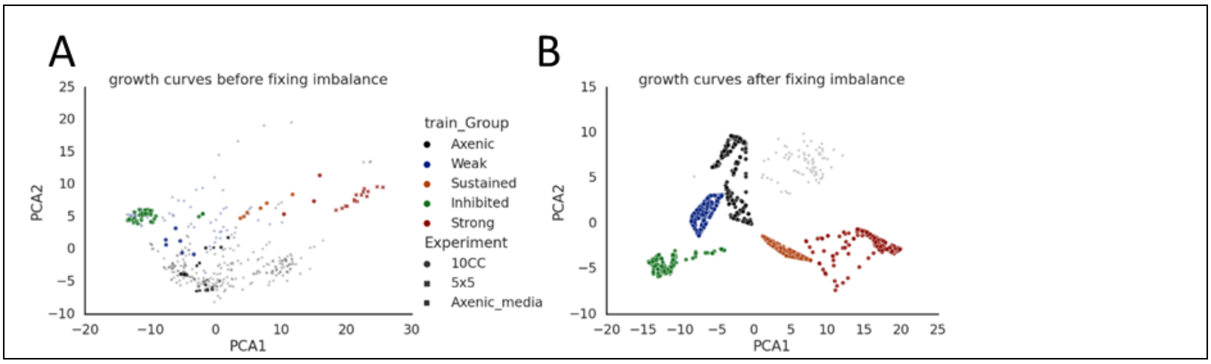

Supplementary Figure S7: Training set of the classification model. A) all growth curves used for training, colored by their class. Shapes indicate source study: 10CC: this study, 5x5: [2] , Axenic_Media: Soussan-Farhat et al, manuscript in prep

We have previously used the growth and decline curves from multiple *Prochlorococcus*-Alteromonas co-cultures to seek similarities between outcomes and identify important time-points [2]. Building upon this approach, we aimed to use both the shapes of the growth curves and important curve features in the classification process. The growth and decline curves themselves were the N biomass and C biomass of *Prochlorococcus*. These curves are linearly correlated in the training set because we assumed a fixed C:N ratio when converting fluorescence to biomass, but they are decoupled in the simulations, which enable a flexible C:N ratio. The C and N curves were log-transformed in order to give higher importance to small dynamic shifts. In parallel, we defined a suite of features describing the curves which emphasize the differences between the phenotypes. These included: max day, max biomass, median, mean and standard deviation of the biomass of days 30-60, mean and standard deviation of the biomass of days 60-80, the last day where biomass was above limit of detection threshold, and D90 (day when the biomass declined by 90%). Each of these features was measured independently for N and C. These 3 datasets (C and N biomass and curve features) were combined in a stacking classifier, combining 2 logistic regression classifiers (separately for N and C biomass) and a random forest classifier based on the generated features (Supplementary Figure S8A).

The logistic regression coefficients were high for the Strong outcome on days 20-40 and 80-90 and for the Sustained outcomes on days 50-60. thus, these outcomes are predicted when *Prochlorococcus* biomass is high on these days (Supplementary Figure S8C). The Weak outcome had negative coefficients on days 20-30, and the inhibited had strong negative coefficients on days 5-10. Low *Prochlorococcus* biomass on these days is key for classification for Weak and Inhibited outcomes respectively. In the random forest classifier, the top features were the median and mean biomass at 30-60 days and max biomass (Supplementary Figure S8B). The model was tested by building on a subset and testing on the rest, and by classifying the samples before imbalance correction, it was also tested by comparison to RMSE and by visual inspection.

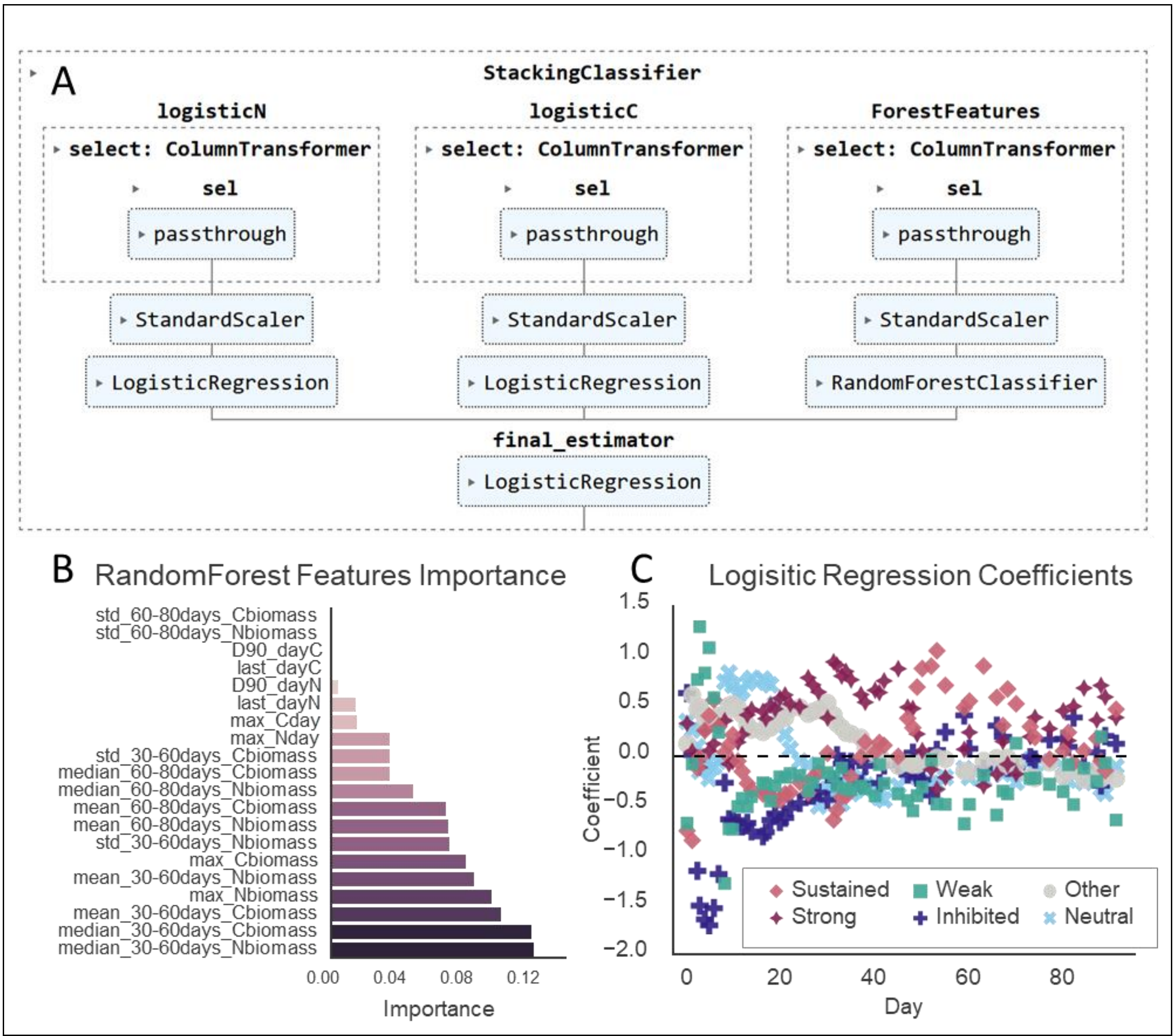


Supplementary Figure S8: The classification stacked model. A) structure of the stacked classification model. 3 estimators: random forest, logistic regression based on vPro N biomass, and logistic regression based on vPro C biomass, are combined via logistic regression final estimator. B) coefficients of logistic regression based on vPro C biomass. C) Random forest features ranked by importance.

We measure the goodness of fit of the classification using both Euclidean distance (RMSE: Root mean square Error) and classification probability (an output of the classification engine) (Supplementary Figure S9). In all models, the majority of the simulations, classified as Weak, had very good fit (RMSE close to 0, and probability close to 1). The fit of Strong and Sustained simulations was not as good, although a higher error is expected for these co-culture outcomes because we are comparing larger biomass expectations and predictions. For Exoenzyme and Overflow simulations, the fit to Strong phenotype had low probability (0.4-0.9) but relatively low error (RMSE 1000-2000), while the fit to Sustained phenotypes was good (high probability and low error). ROS has a good fit to Strong phenotype (high probability and high error), and medium goodness of fit to Sustained (higher error and lower probability). The few simulations that were classified as Inhibited had low probability in all models. Indeed, none of the individual models could recapitulate the complete lack of growth in the inhibited experiments, and this could only occur when some of the models were combined (see main text). The final number of simulations for each outcome is presented in Supplementary Table S4.

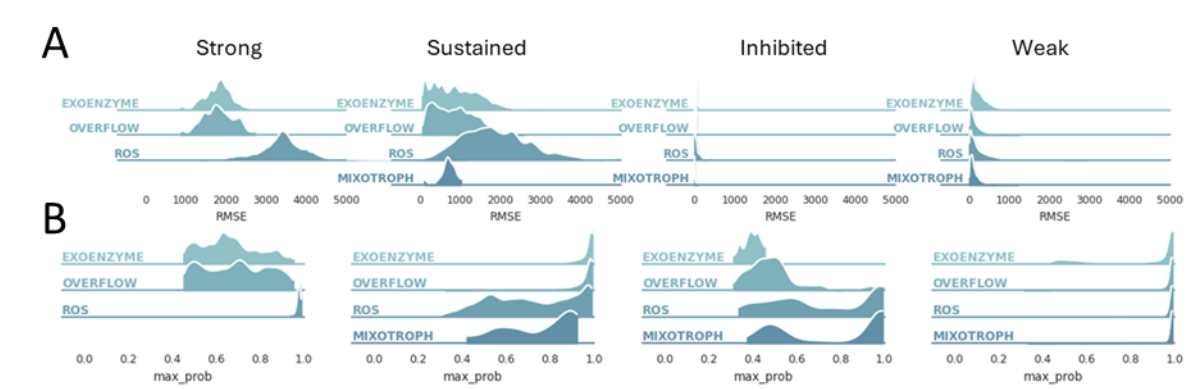


Supplementary Figure S9: Goodness of fit of the coculture classifications. A. Error vs experimental results (RMSE: Root mean square Error). B. Probability of fitting in the stacked ML model

Supplementary Table S4: Number of Montecarlo simulations per phenotype in the different models

| | Versatile vPros | Strong | Sustained | Inhibited | Weak | Neutral | Other |
|---|---|---|---|---|---|---|---|
| EXOENZYME | 21% (135/641) | 1% (622) | 17% (8700) | 0.02% (10) | 20% (10217) | 48% (24513) | 13% (6828) |
| OVERFLOW | 31% (300/970) | 0.2% (99) | 2% (1303) | 0.04% (23) | 49% (29791) | 45% (27817) | 4% (2175) |
| ROS | 2% (11/728) | 2% (547) | 0.1% (19) | 0.1% (17) | 77% (17424) | 16% (3634) | 5% (1030) |
| MIXOTROPH | 0.3% (4/1239) | 0% (0) | 0.03% (12) | 0.2% (56) | 58% (21119) | 42% (15344) | 0.5% (179) |

Our classification scheme was tested by comparison to RMSE. A majority (53%-82%) of the classifications are the same. A sample of the disagreements were inspected manually to verify that the ML predicted outcome is in line with the experimental results.

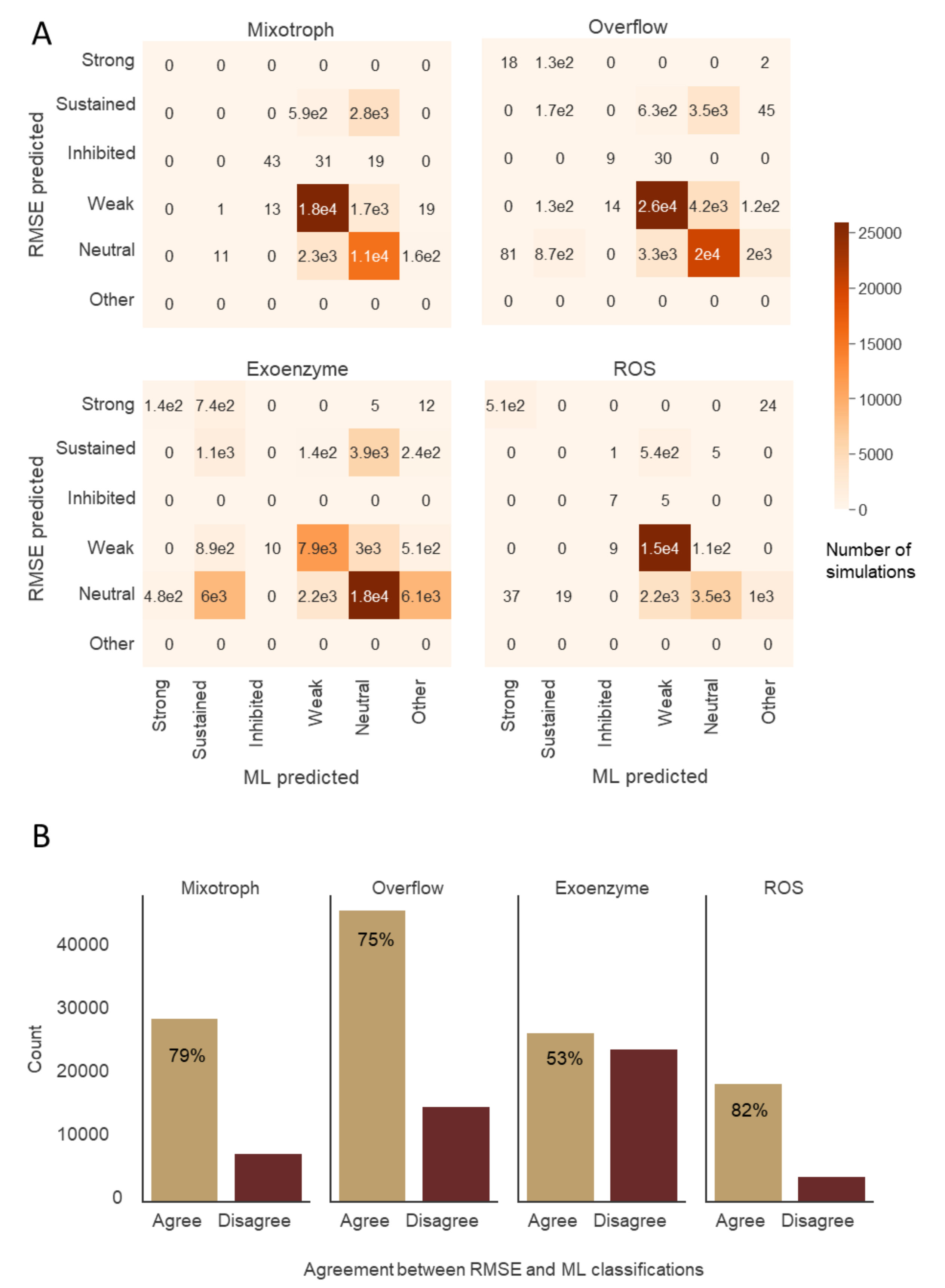


Supplementary Figure S10: Agreement between classification based on RMSE and classification based on stacked ML approach. A. Confusion matrixes per model comparing RMSE-based and ML-based classifications. B. Agreement and disagreements between the classification approaches. Percents on the agreement bars is the % of classifications were both approaches agree.

#### Supplementary Text 7: Estimating the number of *Prochlorococcus* and heterotroph cells in co-culture

It is difficult to estimate the number of *Prochlorococcus* and heterotrophs in the coculture. Flow cytometry can be used to count cells after staining with a DNA dye (e.g. Sybr Green) but is limited because as batch cultures age, the media becomes cluttered with detritus and debris from decomposing dead cells. The culture may also contain aggregates and biofilm as well as *Prochlorococcus* cells, both viable cells and chlorotic (low chlorophyll) which may be non-viable [3]. Finally, as the cultures age, heterotroph cells may shrink due to starvation, whereas some *Prochlorococcus* strains or populations may become larger making it impossible to distinguish between them based on size alone [41].

With this in mind, we used two orthogonal assays to estimate the number of heterotroph cells at the end of the co-culture: flow cytometry (Figure S11D) and Most Probable Number (MPN, Table 1, Supplementary Figure S11D). We counted all of the cells in the samples using flow cytometry and subtracted the number of *Prochlorococcus* cells counted without staining (i.e. based on chlorophyll autofluorescence). These results had a large margin of error, e.g. for the axenic *Prochlorococcus* controls during late stages, which contain no heterotrophs, we identified up to $7x10^6$ unidentified cells. These may represent chlorotic cells or cell debris yet were ~100 fold more abundant than the "healthy" autofluorescent *Prochlorococcus* cells found (Supplementary Figure S11C). To obtain an independent estimate of the size of the heterotroph population, on the last day of the experiment, we employed a second measurement technique, Most Probable Number (MPN), also known as dilution to extinction. In this method, cells are serially diluted and transferred to fresh media, followed by identification of the highest cell dilution (i.e. minimal cell number) where growth was nevertheless observed. MPN has its own caveats, as a statistical method it is inaccurate and can only provide an order-of-magnitude estimate and not an accurate count. It is also limited to culturable microbes which can grow in the test media (e.g. can be affected by bacteria entering into a viable but not culturable state). On the other hand, the assay is orthogonal to flow cytometry and provides additional evidence of the number of viable heterotrophic cells in the culture. The results of the MPN analysis are shown in Supplementary Figures S11D and in Table 1 of the main text.

Combining the results of the flow cytometry and MPN analyses, we identify that, in most cocultures, there are more heterotrophs than *Prochlorococcus* cells at the end of the experiment, with the exception of *Ruegeria pomeroyi*. For most co-cultures, this supports the exoenzyme or overflow models, which both predict a high phototroph-heterotroph ratio (Figure 4A,B). However, in some cases the results may be a bit more nuanced:

1. *Alteromonas* and *Pseudoalteromonas* are both strong phenotypes, and since the flow cytometry and MPN suggest higher numbers of heterotrophs (and higher calculated C and N heterotrophic biomass) this supports either overflow or exoenzyme mechanisms in both these interactions.

2. *Sulfitobacter* and *Ruegeria* are both sustained. The number of heterotrophic cells for Sulfitobacter after 129 days is similar to or higher than *Prochlorococcus*, supporting overflow, exoenzyme or mixotrophy mechanisms. In contrast, the number of *Ruegeria* cells counted by flow cytometry after 129 days are much lower that *Prochlorococcus*, and

no growth at all was observed by MPN (the numbers in the log-log plot in Supplementary Figure S11C represent highest possible abundance). While this could suggest that the ROS mechanism underlies the *Ruegeria-Prochlorococcus* interaction, flow cytometry from earlier time-points suggests higher heterotroph numbers, consistent with overflow, exoenzyme or mixotrophy mechanisms (Supplementary Figure S11D). It is unclear whether the mechanism of interaction changed in this co-culture or whether the mortality rate of the *Ruegeria* cells increases substantially at the late stages of co-culture. Neither of these options is captured by our model formulation.

3. *Marinovum* and *Marinobacter* were both characterized as weak interactions, yet the heterotroph number were much higher than the *Prochlorococcus* ones. This is possibly consistent with exoenzyme, overflow or mixotrophy mechanisms (Figure 4B). We note however that the *Prochlorococcus* numbers at the end of the experiment were very low in these interactions, resulting in a het/Pro ratio much larger than in most models.
4. *Roseovarius* and *Phaeobacter* both inhibited *Prochlorococcus*, a phenotype not captured by most of our individual models – only in model combinations (see main text).

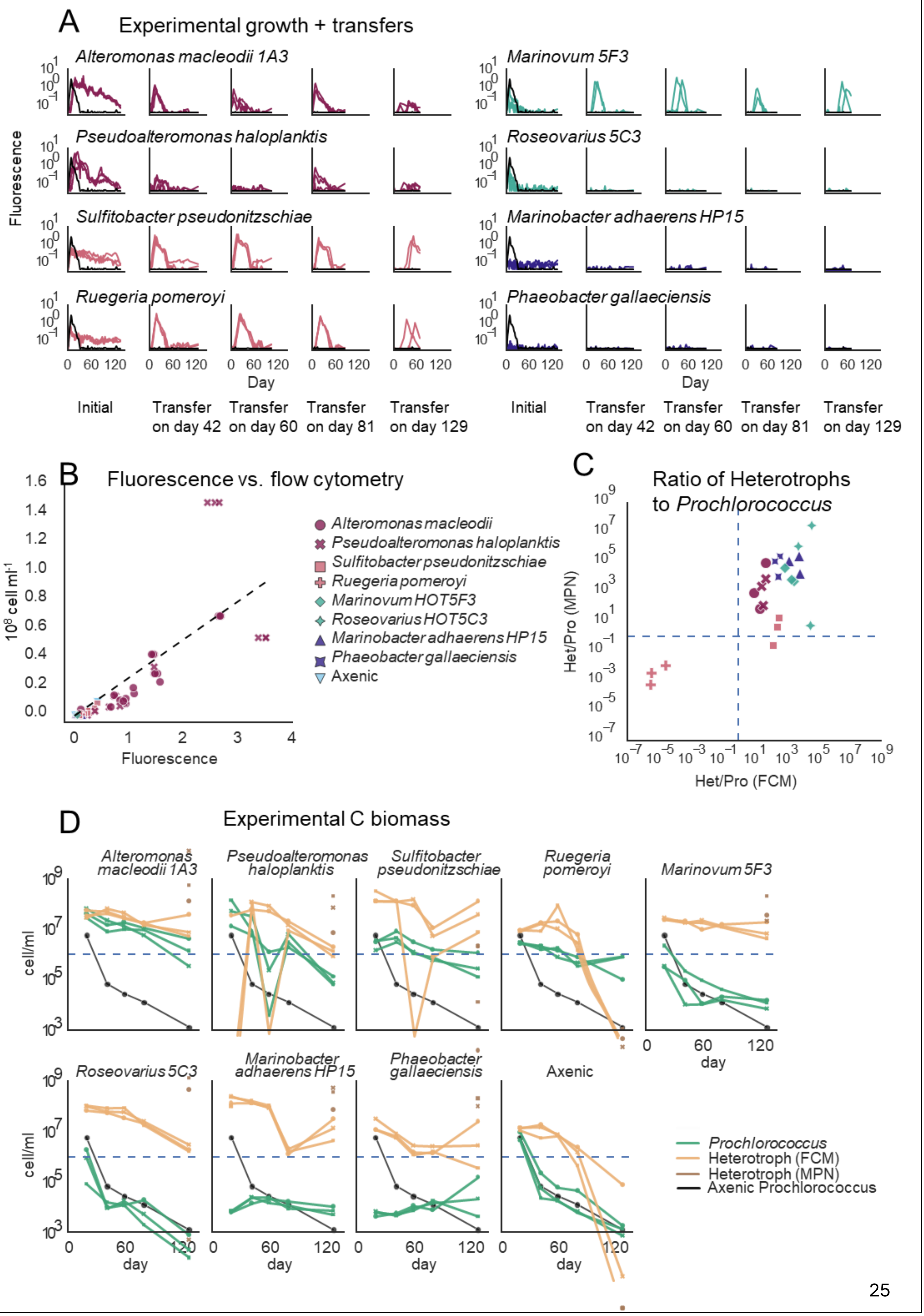

A
Experimental growth + transfers
Alteromonas macleodii 1A3
Pseudoalteromonas haloplanktis
Sulfitobacter pseudonitzschiae
Ruegeria pomeroyi
Marinovum 5F3
Roseovarius 5C3
Marinobacter adhaerens HP15
Phaeobacter gallaeciensis
Fluorescence
Day
Initial
Transfer on day 42
Transfer on day 60
Transfer on day 81
Transfer on day 129
B
Fluorescence vs. flow cytometry
$10^8$ cell ml$^{-1}$
Fluorescence
Alteromonas macleodii
Pseudoalteromonas haloplanktis
Sulfitobacter pseudonitzschiae
Ruegeria pomeroyi
Marinovum HOT5F3
Roseovarius HOT5C3
Marinobacter adhaerens HP15
Phaeobacter gallaeciensis
Axenic
C
Ratio of Heterotrophs to Prochlorococcus
Het/Pro (MPN)
Het/Pro (FCM)
D
Experimental C biomass
Alteromonas macleodii 1A3
Pseudoalteromonas haloplanktis
Sulfitobacter pseudonitzschiae
Ruegeria pomeroyi
Marinovum 5F3
Roseovarius 5C3
Marinobacter adhaerens HP15
Phaeobacter gallaeciensis
Axenic
cell/ml
day
Prochlorococcus
Heterotroph (FCM)
Heterotroph (MPN)
Axenic Prochlorococcus

Supplementary Figure S11: Detailed dynamics of the cuculture growth experiments. A) viability of *Prochlorococcus* as measured by growth after transfer to fresh media. B) Flowrescence vs. Flow cytomrty and fitted linear regression line (adj $R^2$=0.733). C) Ratio of heterotrophs to *Prochlorococcus* on day 129 of the experiment as measured by two independent measures of Heterotroph concentration. D) Experimental C biomass based on Flow cytometry and MPN measurements of *Prochlorococcus* and the heterotrophs during long-term.

#### Supplemental Figures

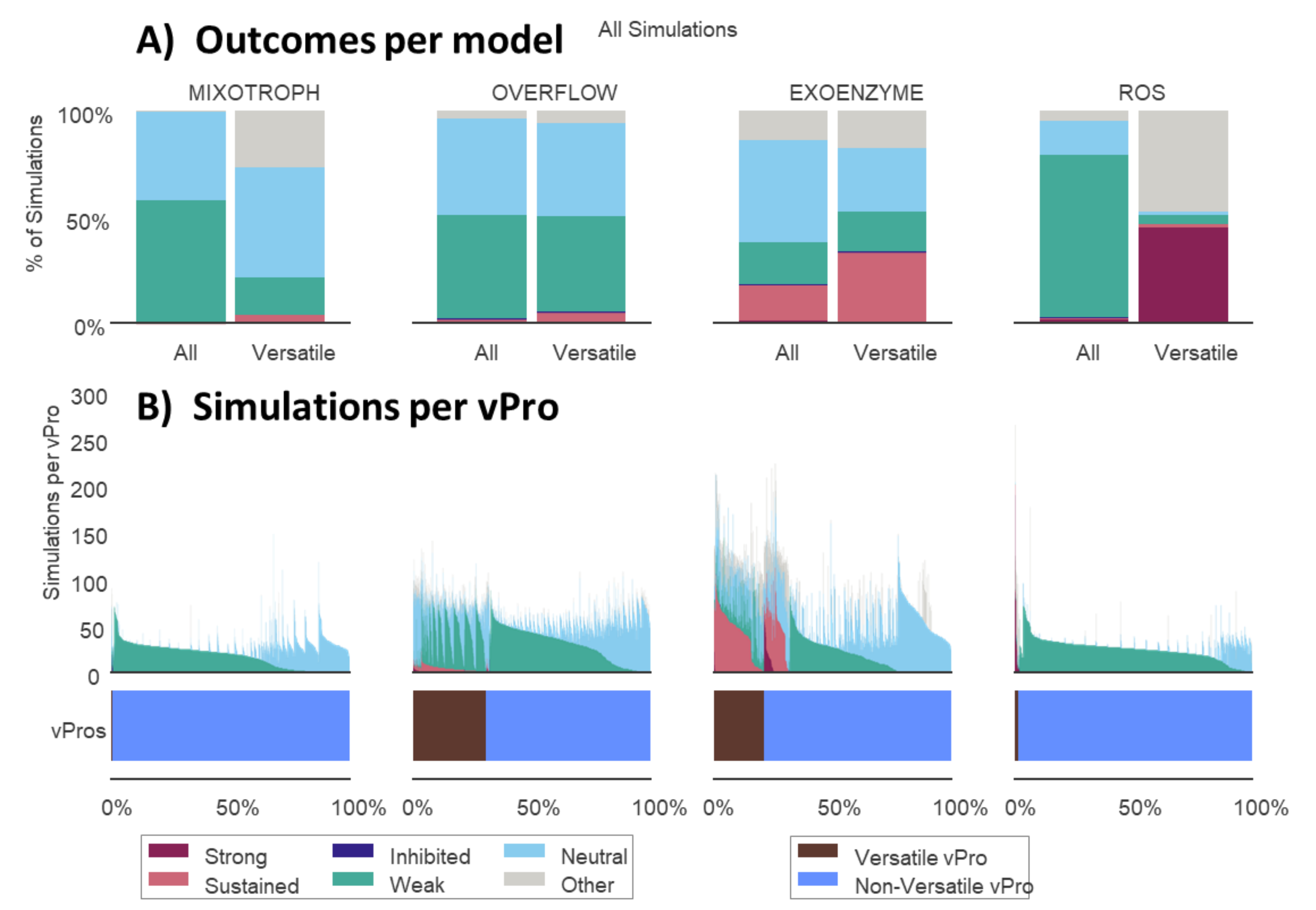


Supplementary Figure S12: outcomes of the simulations used to enlarge the vPro selection from 30-50 vPros to 300-700 vPros. A. Simulation outcomes in each model when either all vPro-vHet pairs are taken into account, or only pairs with a versatile vPro. B. Simulation per vPro, each bar corresponds to simulations of a specific vPro, vPro versatility is indicated at the bottom of the panel.

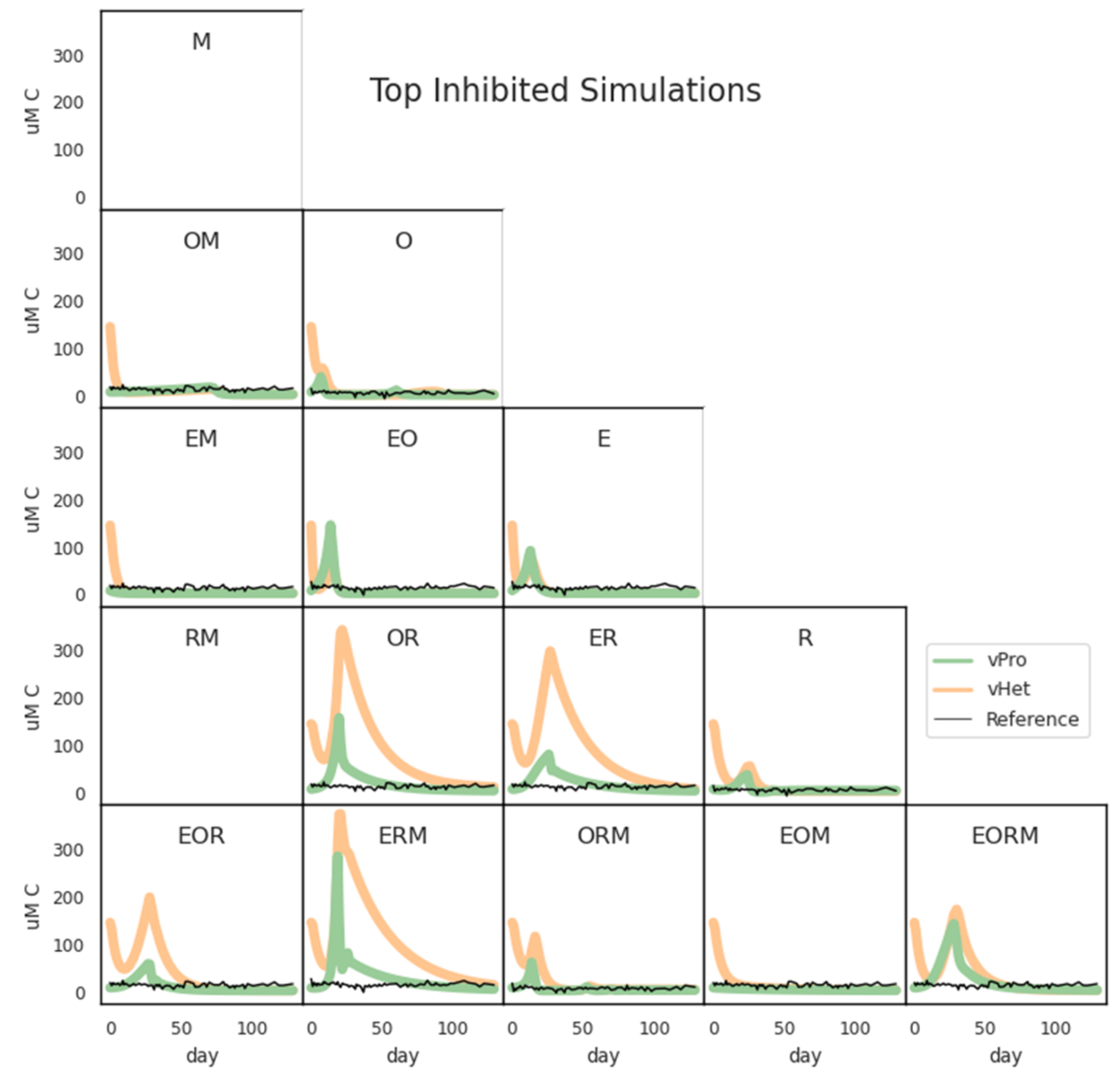


Supplementary Figure S13: Examples of simulations with inhibited outcome across all models combinations. M: Mixotrophy, O: Overflow, E: Exoenzyme, R: ROS. Green line: vPro V biomass, orange line: vHet C biomass, black line: experimental *Prochlorococcus* C biomass (from the coculture experiment).

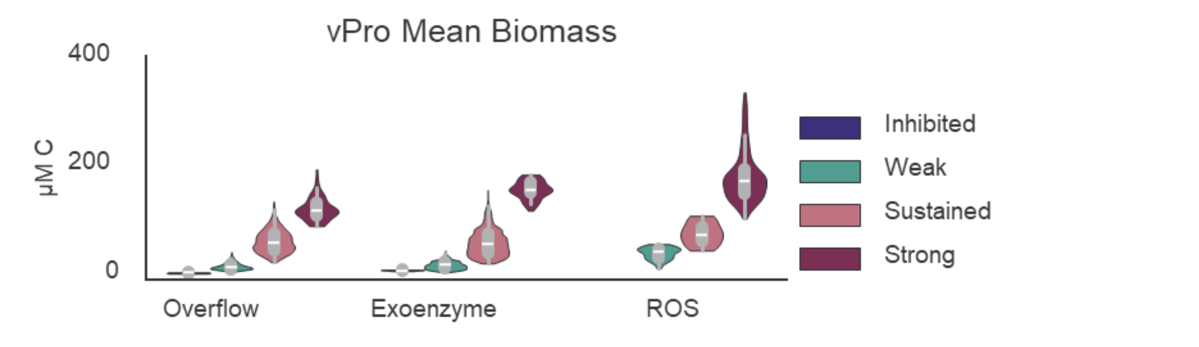


Supplementary Figure S14: Mean C biomass per outcome. In all models there is a positive correlation between the vPro growth and the outcome.

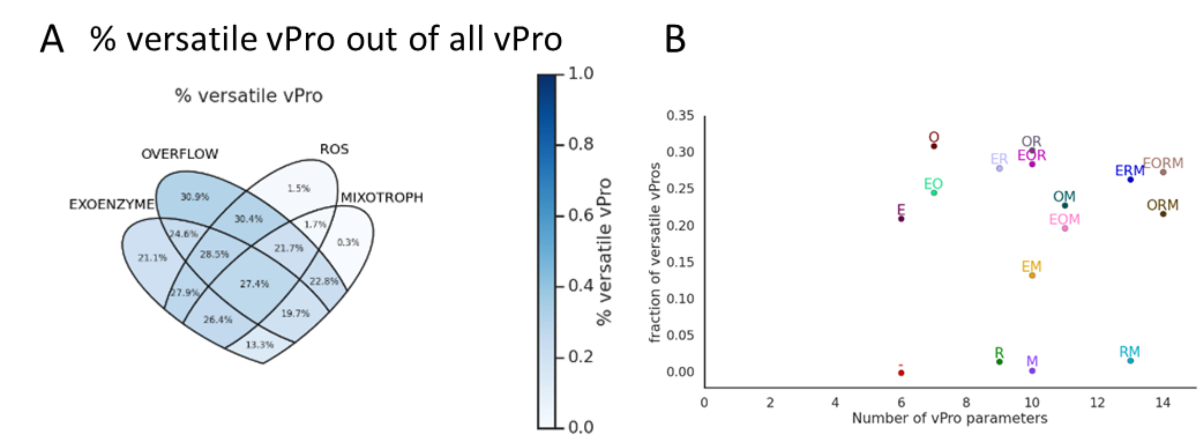


Supplementary Figure S15: Versatile vPros. A) Percent of versatile vPros out of all vPros in all model combinations. B) Percent of versatile vPros out is not correlated with the number of random vPro parameters. M: Mixotrophy, O: Overflow, E: Exoenzyme, R: ROS, -: all mechanism disabled.

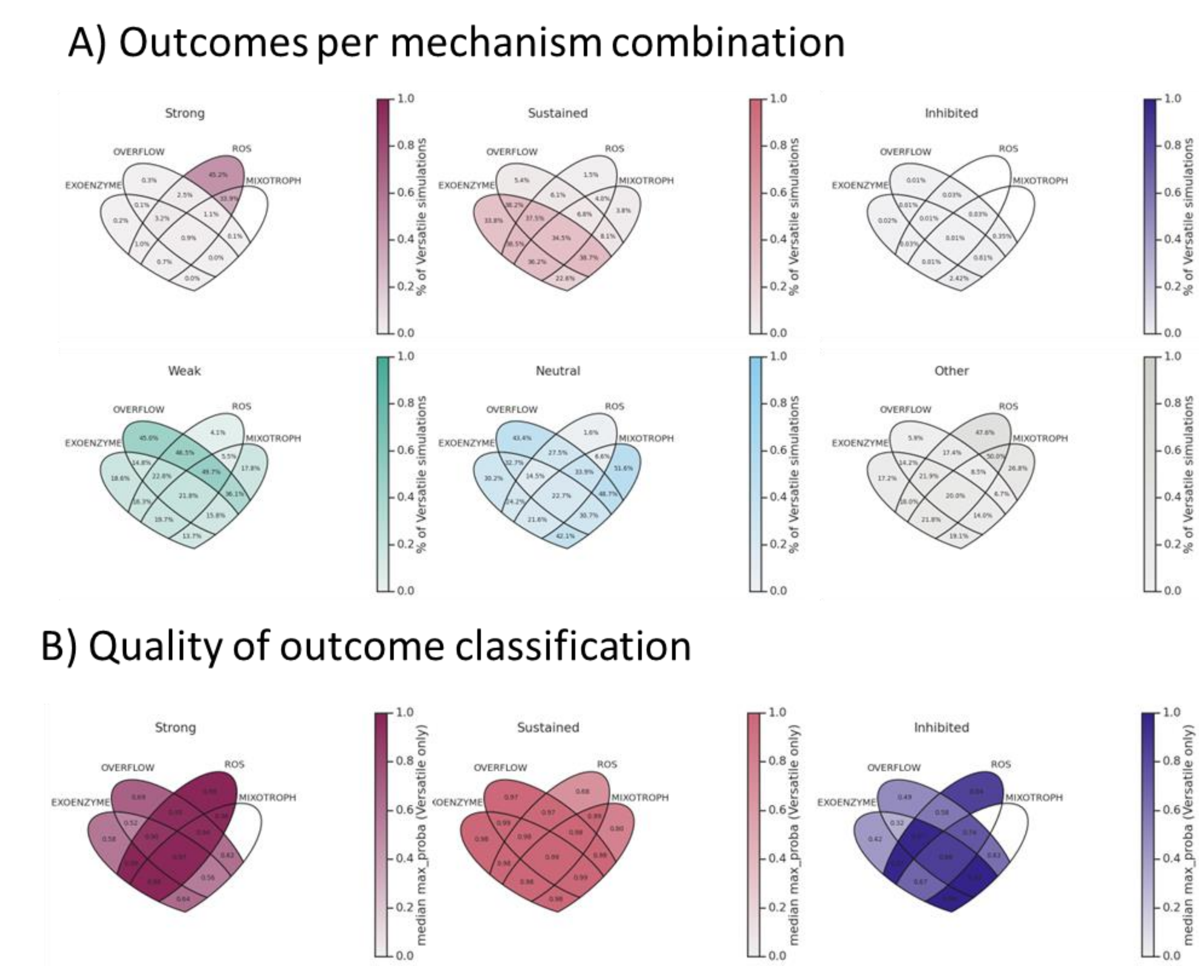


Supplementary Figure S16: outcomes of combining models. A) outcomes of the simulations (percent of simulations with versatile vPros) B) quality of the classification, median of the probability of the classification

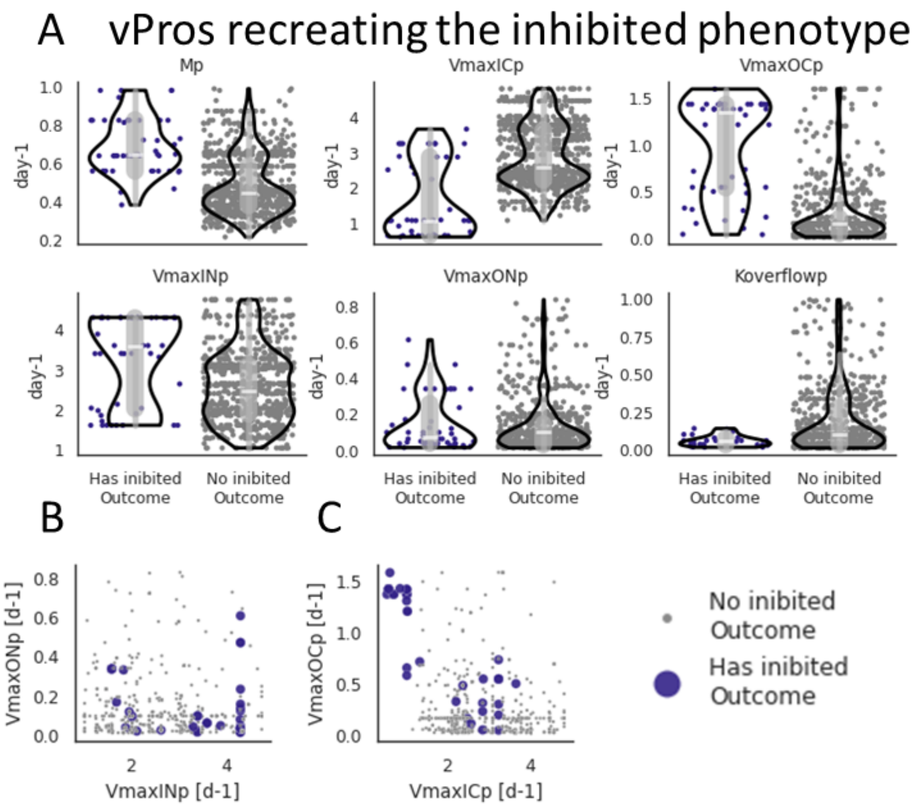


Supplementary Figure S17: parameters of versatile vPros recreating complete *Prochlorococcus* inhibition (models EM, OM, EOM). A) vPro main parameters differentiating vPros that can and cannot recreate inhibition. B) vPro N uptake rates. C) vPro C uptake rates. M: Mixotrophy, O: Overflow, E: Exoenzyme, R: ROS.

543

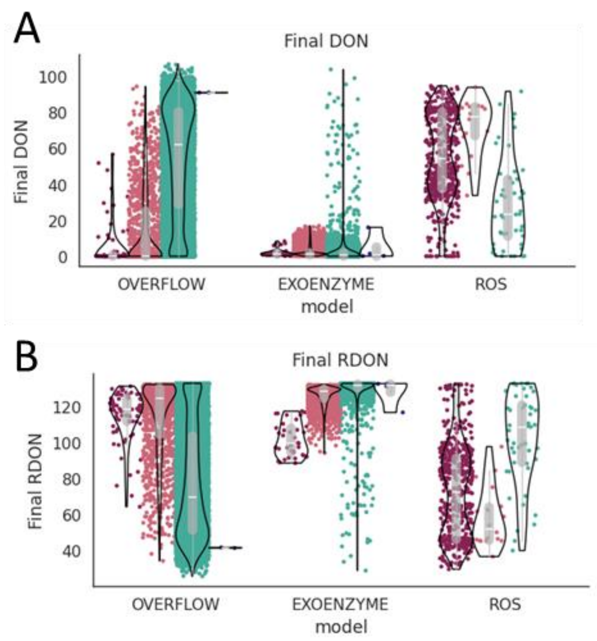


Supplementary Figure S18: N pools at the end of the simulation. A) Final concentration of DON. B) Final concentration of rDON.

544

#### References

1. Aharonovich, D. & Sher, D. Transcriptional response of Prochlorococcus to co-culture with a marine Alteromonas: Differences between strains and the involvement of putative infochemicals. *ISME Journal* **10**, 2892–2906 (2016).

2. Weissberg, O., Aharonovich, D. & Sher, D. Phototroph-heterotroph interactions during growth and long-term starvation across Prochlorococcus and Alteromonas diversity. *ISME Journal* **17**, 227–237 (2023).

3. Roth-Rosenberg, D. *et al.* Prochlorococcus rely on microbial interactions rather than on chlorotic resting stages to survive long-term stress. *mBio* **11**, e01846-20 (2020).

4. Hennon, G. M. M. *et al.* The impact of elevated CO2 on Prochlorococcus and microbial interactions with 'helper' bacterium Alteromonas. *ISME J* **12**, 520 (2017).

5. Morris, J. J., Lenski, R. E. & Zinser, E. R. The black queen hypothesis: Evolution of dependencies through adaptive gene loss. *mBio* **3**, (2012).

6. Biller, S. J., Coe, A. & Chisholm, S. W. Torn apart and reunited: Impact of a heterotroph on the transcriptome of Prochlorococcus. *ISME Journal* **10**, 2831–2843 (2016).

7. Kearney, S. M., Thomas, E., Coe, A. & Chisholm, S. W. Microbial diversity of co-occurring heterotrophs in cultures of marine picocyanobacteria. *Environmental Microbiomes* **16**, (2021).

8. Coe, A. *et al.* Survival of Prochlorococcus in extended darkness. *Limnol Oceanogr* **61**, 1375–1388 (2016).

9. Morris, J. J., Kirkegaard, R., Szul, M. J., Johnson, Z. I. & Zinser, E. R. Facilitation of robust growth of Prochlorococcus colonies and dilute liquid cultures by 'helper' heterotrophic bacteria. *Appl Environ Microbiol* **74**, 4530–4534 (2008).

10. Becker, J. W., Hogle, S. L., Rosendo, K. & Chisholm, S. W. Co-culture and biogeography of Prochlorococcus and SAR11. *ISME J* (2019) doi:10.1038/s41396-019-0365-4.

11. Knight, M. A. & Morris, J. J. Co-culture with Synechococcus facilitates growth of Prochlorococcus under ocean acidification conditions. *Environ Microbiol* **22**, 4876–4889 (2020).

12. Sher, D., Thompson, J. W., Kashtan, N., Croal, L. & Chisholm, S. W. Response of Prochlorococcus ecotypes to co-culture with diverse marine bacteria. *ISME J* **5**, 1125–1132 (2011).

13. Seymour, J. R., Ahmed, T., Durham, W. M. & Stocker, R. Chemotactic response of marine bacteria to the extracellular products of Synechococcus and Prochlorococcus. *Aquatic Microbial Ecology* **59**, 161–168 (2010).

14. Christie-Oleza, J. A., Scanlan, D. J. & Armengaud, J. 'You produce while I clean up', a strategy revealed by exoproteomics during Synechococcus-Roseobacter interactions. *Proteomics* **15**, 3454–3462 (2015).

15. Christie-Oleza, J. A., Sousoni, D., Lloyd, M., Armengaud, J. & Scanlan, D. J. Nutrient recycling facilitates long-term stability of marine microbial phototroph-heterotroph interactions. *Nat Microbiol* **2**, (2017).

16. Kaur, A., Hernandez-Fernaud, J. R., Aguilo-Ferretjans, M. del M., Wellington, E. M. & Christie-Oleza, J. A. 100 Days of marine Synechococcus–Ruegeria pomeroyi interaction: A detailed analysis of the exoproteome. *Environ Microbiol* **20**, 785–799 (2018).

17. Seyedsayamdost, M. R., Case, R. J., Kolter, R. & Clardy, J. The Jekyll-and-Hyde chemistry of phaeobacter gallaeciensis. *Nat Chem* **3**, 331–335 (2011).

18. Töpel, M. *et al.* Complete Genome Sequence of Novel Sulfitobacter pseudonitzschiae Strain SMR1, Isolated from a Culture of the Marine Diatom Skeletonema marinoi . *J Genomics* **7**, 7–10 (2019).

19. Amin, S. A. *et al.* Interaction and signalling between a cosmopolitan phytoplankton and associated bacteria. *Nature* **522**, 98–101 (2015).

20. Gärdes, A., Ramaye, Y., Grossart, H. P., Passow, U. & Ullrich, M. S. Effects of Marinobacter adhaerens HP15 on polymer exudation by Thalassiosira weissflogii at different N:P ratios. *Mar Ecol Prog Ser* **461**, 1–14 (2012).

21. Clark, D. R., Rees, A. P. & Joint, I. *Ammonium Regeneration and Nitrification Rates in the Oligotrophic Atlantic Ocean: Implications for New Production Estimates*. vol. 53 (2008).

22. Wu, Z. *et al.* Single-cell measurements and modelling reveal substantial organic carbon acquisition by Prochlorococcus. *Nat Microbiol* **7**, 2068–2077 (2022).

23. Cole, J. J. & Caraco, N. F. Atmospheric exchange of carbon dioxide in a low-wind oligotrophic lake measured by the addition of SF6. *Limnol Oceanogr* **43**, 647–656 (1998).

24. Williams, R. G. & Follows, M. J. *Ocean Dynamics and the Carbon Cycle: Principles and Mechanisms*. (Cambridge University Press, 2011).

25. Redfield, A. C. On the Proportions of Organic Derivatives in Sea Water and Their Relation to the Composition of Plankton. in *James Johnstone Memorial Volume* 176–192 (University Press of Liverpool, 1934).

26. J. Geider, R. & Osborne, B. A. Respiration and microalgal growth: a review of the quantitative relationship between dark respiration and growth. *New phytologist* **112**, 327–341 (1989).

27. Follows, M. J., Dutkiewicz, S., Grant, S. & Chisholm, S. W. Emergent biogeography of microbial communities in a model ocean. *Science (1979)* **315**, 1843–1846 (2007).

28. Muñoz-Marín, M. del C., López-Lozano, A., Moreno-Cabezuelo, J. Á., Díez, J. & García-Fernández, J. M. Mixotrophy in cyanobacteria. *Curr Opin Microbiol* **78**, 102432 (2024).

29. Coe, A. *et al.* Coping with darkness: The adaptive response of marine picocyanobacteria to repeated light energy deprivation. *Limnol Oceanogr* **66**, 3300–3312 (2021).

30. Ofaim, S., Sulheim, S., Almaas, E., Sher, D. & Segrè, D. Dynamic Allocation of Carbon Storage and Nutrient-Dependent Exudation in a Revised Genome-Scale Model of Prochlorococcus. *Front Genet* **12**, (2021).

31. Dubinsky, Z. & Berman-Frank, I. Uncoupling primary production from population growth in photosynthesizing organisms in aquatic ecosystems. *Aquat Sci* **63**, 4–17 (2001).

32. Azam, F. & Malfatti, F. Microbial structuring of marine ecosystems. *Nat Rev Microbiol* **5**, 966-U23 (2007).

33. Ma, L., Calfee, B. C., Morris, J. J., Johnson, Z. I. & Zinser, E. R. Degradation of hydrogen peroxide at the ocean's surface: the influence of the microbial community on the realized thermal niche of Prochlorococcus. *ISME J* **12**, 473 (2017).

34. Morris, J. J., Johnson, Z. I., Szul, M. J., Keller, M. & Zinser, E. R. Dependence of the cyanobacterium Prochlorococcus on hydrogen peroxide scavenging microbes for growth at the ocean's surface. *PLoS One* **6**, e16805 (2011).

35. Grossowicz, M. *et al.* Prochlorococcus in the lab and in silico: The importance of representing exudation. *Limnol Oceanogr* **62**, 818–835 (2017).

36. Berthelot, H., Duhamel, S., L'Helguen, S., Maguer, J. F. & Cassar, N. Inorganic and organic carbon and nitrogen uptake strategies of picoplankton groups in the northwestern Atlantic Ocean. *Limnol Oceanogr* **66**, 3682–3696 (2021).

37. Masuda, T. *et al.* Coexistence of Dominant Marine Phytoplankton Sustained by Nutrient Specialization. *Microbiol Spectr* **11**, e04000-22 (2023).

38. Maranon, E. *et al.* Unimodal size scaling of phytoplankton growth and the size dependence of nutrient uptake and use. *Ecol Lett* **16**, 371–379 (2013).

39. Ayo, B. *et al.* Kinetics of glucose and amino acid uptake by attached and free-living marine bacteria in oligotrophic waters. *Mar Biol* **138**, 1071–1076 (2001).

40. Muñoz-Marín, M. del C. *et al.* Glucose uptake in Prochlorococcus: Diversity of kinetics and effects on the metabolism. *Front Microbiol* **8**, (2017).

41. Roth-Rosenberg, D., Aharonovich, D., Omta, A. W., Follows, M. J. & Sher, D. Dynamic macromolecular composition and high exudation rates in Prochlorococcus. *Limnol Oceanogr* **66**, 1759–1773 (2021).

42. Hopkinson, B. M., Young, J. N., Tansik, A. L. & Binder, B. J. The minimal CO2-concentrating mechanism of prochlorococcus spp. MED4 is effective and efficient. *Plant Physiol* **166**, 2205–2217 (2014).

43. Muñoz-Marín, M. del C. *et al.* Differential Timing for Glucose Assimilation in Prochlorococcus and Coexistent Microbial Populations in the North Pacific Subtropical Gyre . *Microbiol Spectr* **10**, (2022).

44. Stepanauskas, R. Differential Dissolved Organic Nitrogen Availability and Bacterial Aminopeptidase Activity in Limnic and Marine Waters. *Microb Ecol* **38**, 264–272 (1999).

45. Sutherland, K. M. *et al.* Extracellular superoxide production by key microbes in the global ocean. *Limnol Oceanogr* **64**, 2679–2693 (2019).

46. Bond, R. J., Hansel, C. M. & Voelker, B. M. Heterotrophic Bacteria Exhibit a Wide Range of Rates of Extracellular Production and Decay of Hydrogen Peroxide. *Front Mar Sci* **7**, (2020).

47. Roe, K. L., Schneider, R. J., Hansel, C. M. & Voelker, B. M. Measurement of dark, particle-generated superoxide and hydrogen peroxide production and decay in the subtropical and temperate North Pacific Ocean. *Deep Sea Res 1 Oceanogr Res Pap* **107**, 59–69 (2016).
